# Global invasion history and genomic signatures of adaptation of a highly invasive lace bug

**DOI:** 10.1101/2024.03.26.586866

**Authors:** Zhenyong Du, Xuan Wang, Yuange Duan, Shanlin Liu, Li Tian, Fan Song, Wanzhi Cai, Hu Li

## Abstract

Invasive species cause enormous economic and ecological damage. Climate change has resulted in an unprecedented increase in the number and impact of invaders. The mechanisms underlying invasions, however, are largely unclear. The sycamore lace bug, *Corythucha ciliata*, is a highly invasive species that originated in North America. Its population has expanded over the Northern Hemisphere since the 1960s. In this study, we assemble the lace bug’s genome using high-coverage PacBio, Illumina, and Hi-C sequencing. We identify 15,278 protein-coding genes and expansion of gene families with oxidoreductase and metabolic activities. In-depth resequencing of 402 samples collected from native and nine invasive countries across three continents identified 2.74 million single nucleotide polymorphisms. We resolve two major invasion routes of this lace bug from North America through both Europe and Japan, forming a contact zone in East Asia. Genomic signatures of selection associated with invasion and long-term balancing selection in native ranges are identified. These genomic signatures overlap with each other and the expanded genes, suggesting an improvement in the oxidative stress and thermal tolerance of this species. Our findings offer valuable insights into the genomic architecture and adaptive evolution underlying the invasive capabilities of species in the face of rapid environmental changes.

## Introduction

Biological invasions threaten ecosystems, biodiversity, public health, and welfare, leading to enormous economic losses [1,2]. Global climate change and rising temperatures have resulted in an unprecedented rise in the number and impact of invaders, with invasive ranges surpassing native ranges [3–7]. Understanding the mechanisms underlying successful invasions is an important goal, as only a small proportion of introduced species become effective invaders [8,9]. Various hypotheses based on transport opportunities, propagule pressure, habitat matching, fecundity, and population size have been suggested [10–12]; however, consistent empirical evidence across invasion scenarios and organisms is lacking. Few studies have investigated the genomic signatures underlying this variation.

Both pre- and post-introduction adaptations critically contribute to successful invasion and establishment. Given that the environmental differences between introduced and native ranges might result in strong selective pressures, genetic adaptation post-introduction to novel habitats is necessary for colonization and invasion [12–14]. Natural selection favors only a few fitness-enhancing genes [15–17], and evolution rates could be quite high during invasion [18–20]. In addition, an invasive species might be pre-adapted to part or most of the novel environment. Phenotypic plasticity may be sufficient for invaders to survive and establish invasive ranges, particularly if the native and invasive ranges are highly similar [12]. The potential for pre-introduction invasive capacity may be related to the species’ genomic architecture [12,13,21,22], and breadth of their native distribution that are related to fluctuating environments favoring more alleles at a given locus to maintain functional polymorphisms [6,8,12,19,23,24].

The sycamore lace bug, *Corythucha ciliata* (Heteroptera: Tingidae), is an oligophagous grazer specialized to sycamore (*Platanus* spp.) trees (Figure 1B). *Corythucha ciliata* nymphs and adults feed on the underside of leaves, resulting in a white stippling [25], which subsequently leads to chlorotic or bronzed leaf foliage, tree growth inhibition, premature senescence, and even tree death. The sycamore lace bug is native to North America [25,26]. Invasive populations were first discovered in Padova, Italy, in 1964 and subsequently recorded in additional Southern and Central European countries. Between 1995–2002, *C. ciliata* also invaded South Korea, China, and Japan [27–30]. Our sampling during 2016–2019 (Table S1) revelated its distribution range in East Asia has had spread to most cities in southwestern, northeastern, eastern, and central China [31] as well as most cities in South Korea and to Honshu, Shikoku, and Kyushu in Japan. The lace bug has also been discovered in Chile [32], Australia, and South Africa [33]. As sycamore trees being popular street trees worldwide, widespread invasion of sycamore lace bug populations has severely damaged urban afforestation. Despite the constant rise in temperature owing to climate change, this temperate species has also extensively invaded multiple subtropical regions [31]. However, the global invasion routes and the mechanisms underlying both the rapid invasion and the variation between invasive and native ranges are unclear.

**Figure 1.**
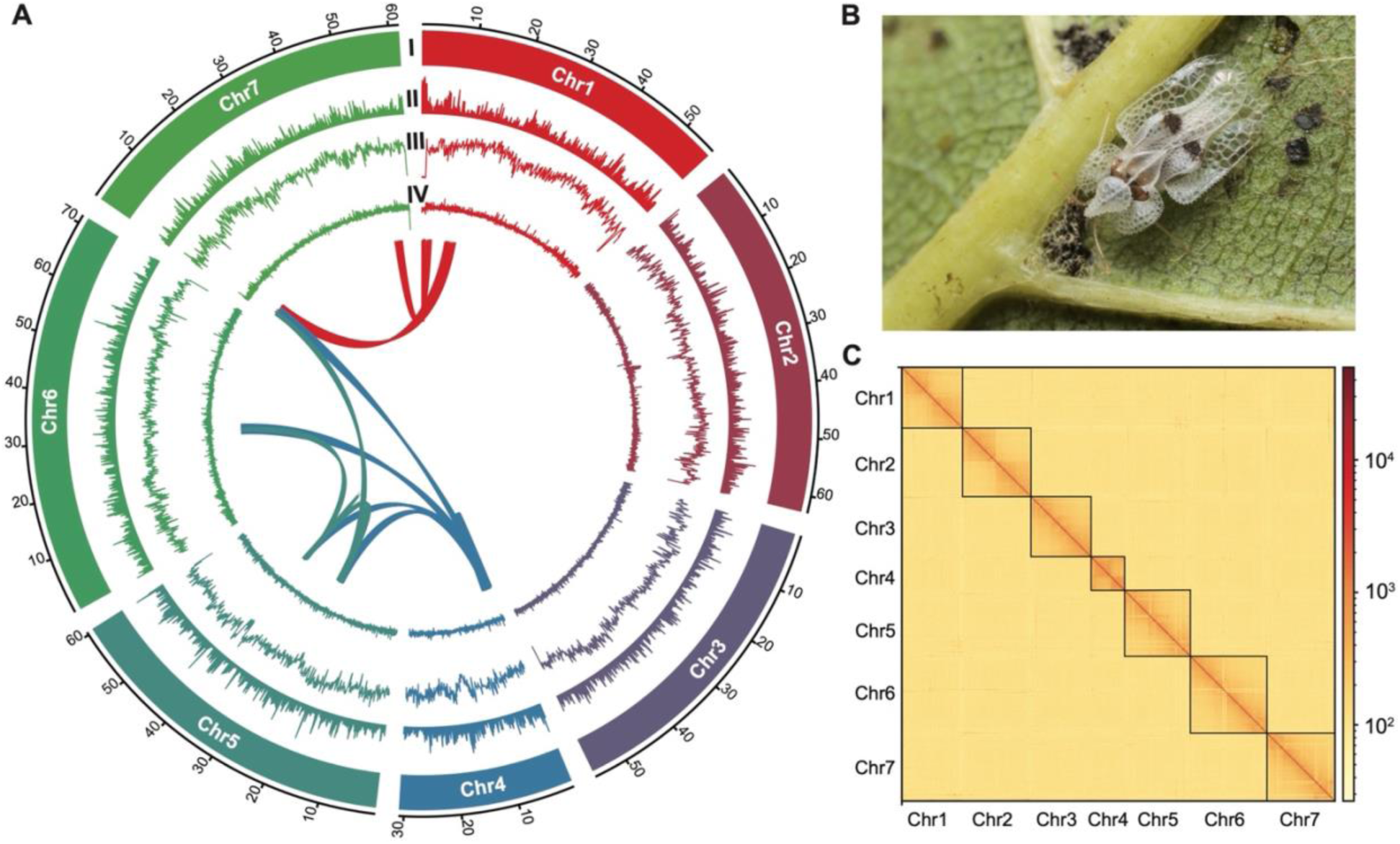
Genome architecture of *Corythucha ciliata*. **A.** Circular diagram exhibiting the landscape of the *C. ciliata* genome. I. Seven chromosomes on the Mb scale. II. Density of protein coding genes. III. Distribution of repeat sequences. IV. Distribution of GC content. The distributions of protein coding genes and repeat sequences are calculated in a 100-kb non-overlapping sliding window. The distribution of GC content is calculated in a 10-kb window. The ribbon inside the circle represents the synteny block within the genome. **B.** *C. ciliata* feeding on a sycamore leaf. **C.** Hi-C contact map of seven chromosomes.

Successful invasion often hinges on both pre- and post-introduction adaptations, yet the mechanisms driving this success remain obscure. We propose that certain genomic features, particularly the expansion of specific gene families and the selection for pivotal traits, might be key to invasiveness. To substantiate our hypothesis, we used *C. ciliata* as a model invasive organism. Our objectives were to (1) pinpoint species-specific gene families that have undergone expansion across the genome; (2) decipher genomic signatures of selection during different invasion scenarios; (3) investigate signatures of selection within the native habitats; and (4) assess the interplay between these adaptive genomic features. By addressing these aims, we hope to unravel the genomic architecture that facilitates the invasive prowess of species like *C. ciliata*.

## Results

### Chromosome-level genome assembly

We used PacBio long-read sequencing and assembly, Illumina short-read polishing, and Hi-C scaffolding to assemble the *C. ciliata* genome. We obtained 9,158,256 PacBio reads, corresponding to 120.03 gigabases (Gb) of data and 302-fold (×) genome coverage, using high-molecular-weight (HMW) DNA extracted from *C. ciliata* samples. Based on k-mer analysis, a haploid genome size of ∼400 Mb was estimated (Figure S1). The initial assembly had 1085 contigs, totaling 400.78 megabases (Mb), and a contig N50 of 1.19 Mb. This assembly was further scaffolded based on 61.62 Gb Hi-C data (∼154×), resulting in a 401.27 Mb final assembly with a scaffold N50 of 60.52 Mb. We anchored 99.67% of the assembly, equivalent to 399.93 Mb, onto seven chromosomes, consistent with findings from previous karyotype studies [34,35] (Figures 1A and 1C; Table S2). Benchmark of Universal Single-Copy Orthologs (BUSCO) analyses indicated that the *C. ciliata* genome captured 95.3% (93.7% single-copy and 1.6% duplicated) of complete BUSCOs (1579 out of 1658 in insecta odb10).

### Genomic annotation and comparative analyses

We identified 15,278 protein-coding genes based on a full-length transcriptome, homologous proteins, and *ab initio* prediction, covering 206.29 Mb in length (Table S3). Of these genes, 13,235 (86.63%) were functionally annotated using five databases (Figures S2 and S3). To perform comparative analyses, we identified orthologous groups (aka. gene families) across ten hemipteran insects, considering the thrips *Frankliniella occidentalis* as an outgroup. We reconstructed a phylogenetic tree using a matrix of 895 concatenated single-copy orthologous genes that supported the phylogenetic framework of Hemiptera ((Auchenorrhyncha + Heteroptera) + Sternorrhyncha) (Figure 2A). We identified a substantial number (5736) of shared gene families among four Hemiptera and Paraneoptera species (Figure 2B).

**Figure 2.**
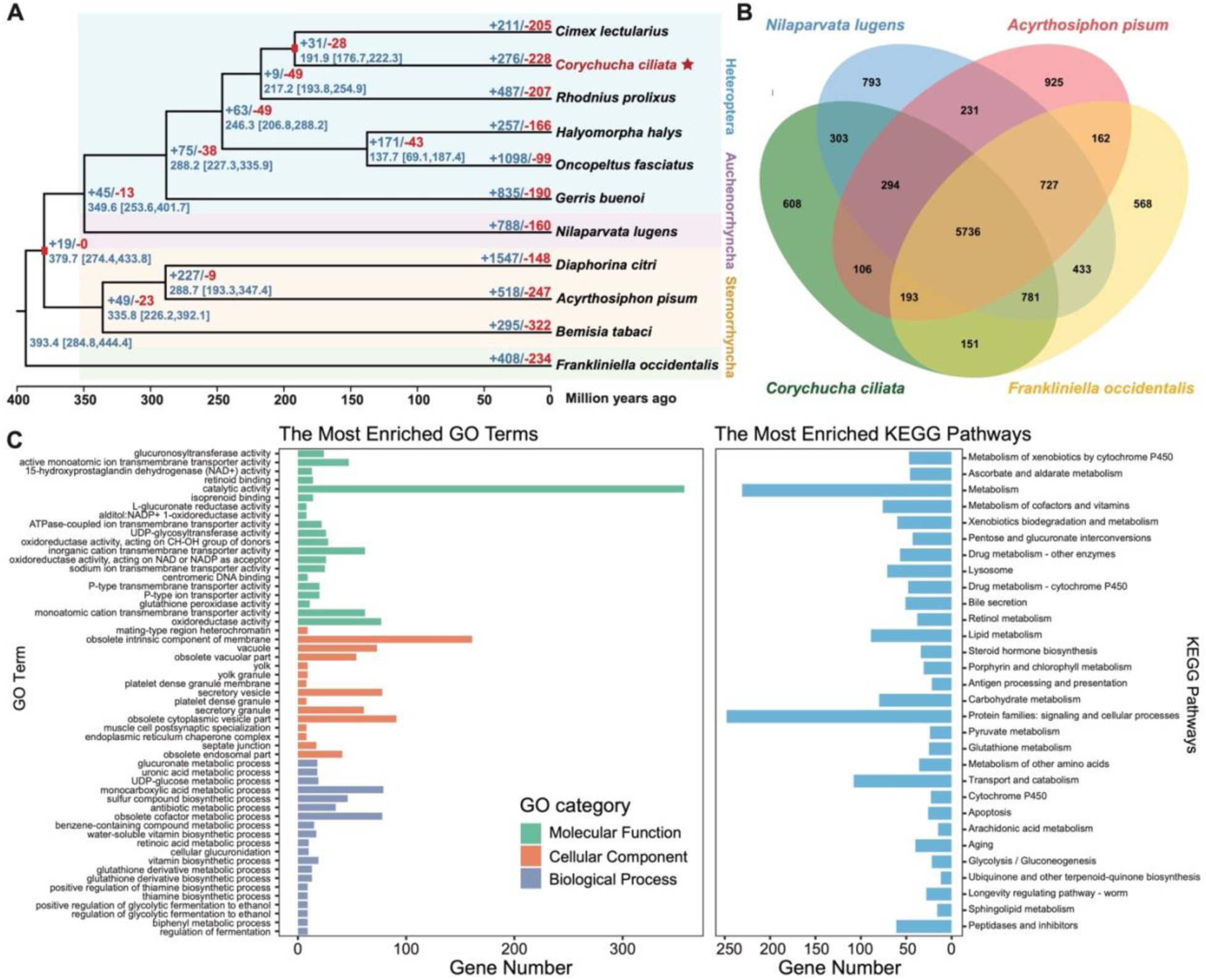
Comparative genomics of hemipteran insects. **A.** Phylogeny, divergence time, and gene family expansion and contraction of ten hemipterans with the thrips *Frankliniella occidentalis* as an outgroup. The phylogenetic relationship tree was reconstructed using RAxML. All nodes show 100% bootstrap support. Divergence time is estimated using MCMCTree, with two calibrated nodes marked with red rectangles. The mean divergence time is shown at each node, with the bracket indicating the 95% highest posterior densities. The numbers of expanded (in blue) and contracted (in red) gene families are indicated on the branch next to each node. **B.** Venn diagram showing the numbers of gene families shared among four paraneopteran species. **C.** GO and KEGG enrichments of the expanded genes in the *C. ciliata* genome. GO terms and KEGG pathways are sorted according to the P value (higher P values toward the bottom). Only the top 20 GO terms of Biological Process and Molecular Function categories, the top 15 GO terms of Cellular Component category, and the top 30 KEGG pathways are shown. The complete list of enriched GO terms and KEGG pathways is given in Table S5 and S6.

To examine the genomic mechanism of *C. ciliata* adaptive capacity, we conducted gene family expansion and contraction analysis across the hemipteran (paraneopteran) phylogeny. We identified 276 expanded gene families corresponding to 1171 genes, and 228 contracted gene families corresponding to 402 genes in the *C. ciliata* genome (Table S4). We performed gene ontology (GO) and Kyoto Encyclopedia of Genes and Genomes (KEGG) enrichment analyses to reveal the potential functions of these genes. The expanded genes were significantly enriched for oxidoreductase and metabolism-related functions. In the Molecular Function (MF) category, the most significantly enriched GO terms included “15-hydroxyprostaglandin dehydrogenase (NAD+) activity” [GO:0016404], “L-glucuronate reductase activity” [GO:0047939], “oxidoreductase activity” [GO:0016491], “alditol:NADP^+^ 1-oxidoreductase activity” [GO:0004032], and “glutathione peroxidase activity” [GO:0004602] (Figure 2C; Table S5). The most significantly enriched Biological Process (BP) GO terms were “glucuronate metabolic process” [GO:0019585], “uronic acid metabolic process” [GO:0006063], and “UDP-glucose metabolic process” [GO:0006011] (Figure 2C; Table S5). “Catalytic activity” [GO:0003824] and “obsolete intrinsic component of membrane” [GO:0031224] were the GO terms with the greatest number of enriched genes. Among the KEGG pathways, “metabolism of xenobiotics by cytochrome P450”, “ascorbate and aldarate metabolism”, and “metabolism” were the most significantly enriched pathways. Overall, the categories of “metabolism”, “protein families: signaling and cellular processes”, and “transport and catabolism” possessed the greatest number of enriched pathways (Figure 2C; Table S6). In comparison, the contracted genes were significantly enriched for sterol, cholesterol, and lipid transfer functions, and for development and morphogenesis. The most significantly enriched GO terms included “ABC-type sterol transporter activity” [GO:0034041], “cholesterol transfer activity” [GO:0120020], and “lipid transfer activity” [GO:0120013] in the MP category, and “ecdysteroid metabolic process” [GO:0045455] and “apoptotic process involved in morphogenesis” [GO:0060561] in the BP category (Tables S7 and S8).

### Genome resequencing and genomic variations

In total, 410 *C. ciliata* samples were sequenced, of which eight were removed due to low mapping rates (Table S9). Four of these eight samples remained available for mitochondrial genome assembly. The remaining 402 samples were used for single nucleotide polymorphism (SNP) genotyping, with coverage depths in the range of 9.03–42.9× and an average of 22.12× (Table S9). Mapping rates to the reference genome ranged from 66.61% to 97.68%, with an average of 95.84% (Table S9). Thirteen samples were excluded due to high missing rates in the SNP dataset (Table S10). Nineteen individuals were identified as full-siblings of another individual within the population and excluded based on the kinship coefficient (Table S11). A total of 2.74 million SNPs from 370 samples progressed for further analysis.

To account for linkage disequilibrium (LD) and avoid bias in the population genetic structure, we filtered and pruned the SNPs to maintain a single SNP per 3-kb window using PLINK, resulting in 106,761 independent loci that were used to analyze population genetic structure.

To better resolve the evolutionary history of *C. ciliata* and integrate the advantages of the haploid organelle genome reflecting the recent divergence history [36–38], a mitochondrial dataset consisting of 665 whole mitochondrial genomes (mitogenomes) was gathered. These mitogenomes included 406 samples sequenced in the present study and 259 obtained from our previous skim sequencing data [37,38] (Table S1). This dataset represents a powerful resource for resolving the global invasion history and adaptation of *C. ciliata*.

### Population genetic structure

ADMIXTURE analysis revealed that all populations possessed six ancestry components (K = 6) based on the lowest error of cross-validation among K = 2–10 (Figures 3B and S4). The native American populations possessed four major components. The European populations inherited one of the four components. The Japanese, South Korean, and Chinese populations exhibited high levels of admixture. Chinese populations shared components with European, South Korean, and Japanese populations. The Japanese Honshu population possessed three components, while the Shikoku and Kyushu populations had one. The South Korean and Chinese populations shared two components (Figure 3B).

**Figure 3.**
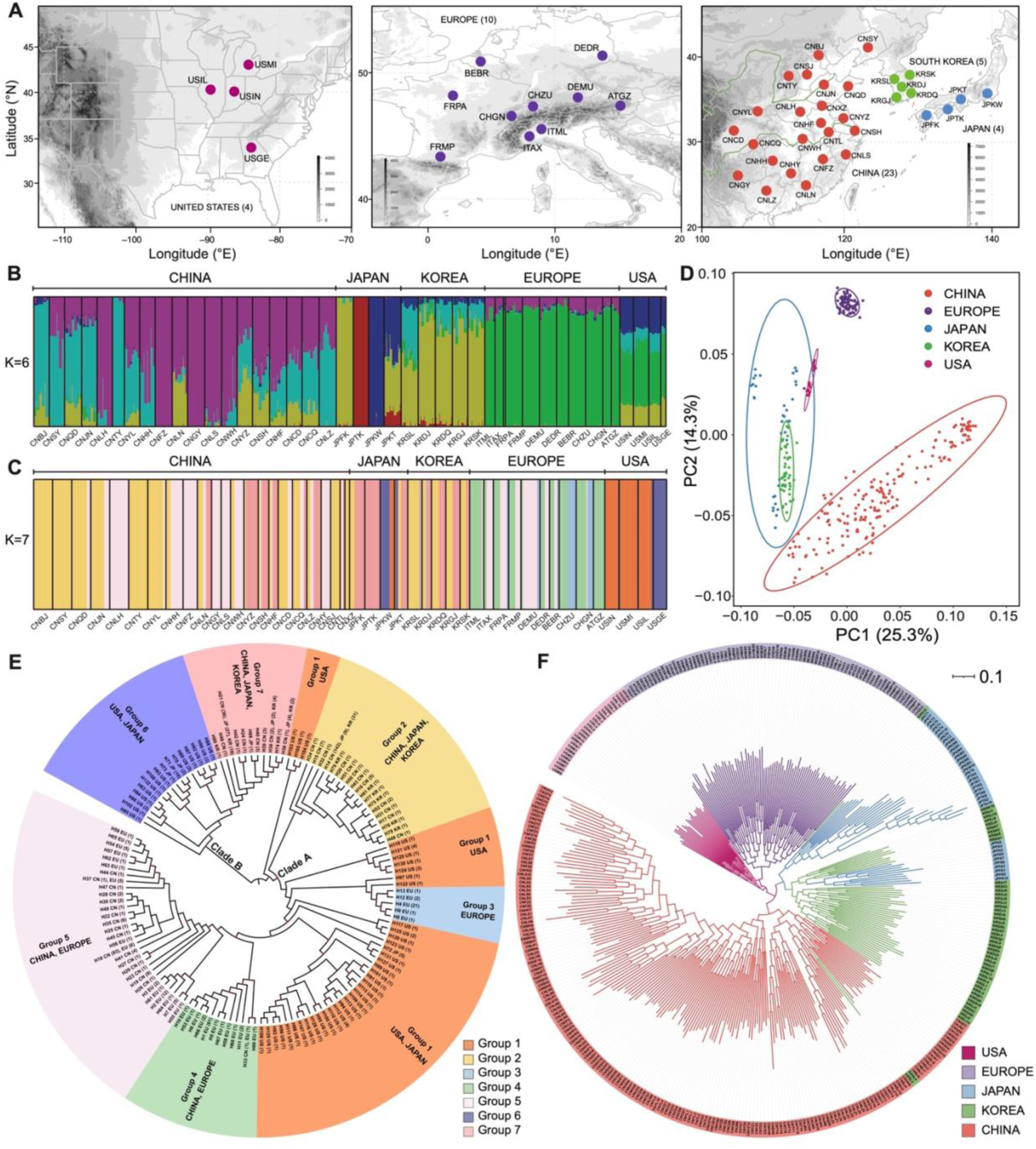
Population sampling and genetic structure based on genomic SNPs and mitogenomes. **A.** Sample localities in North America, Europe, and East Asia. Different colors indicate sample localities in five different regions, i.e., the USA (in amaranth), Europe (in purple), Japan (in blue), South Korea (in green), and China (in orange). **B.** Genetic structure revealed by ADMIXTURE based on genomic SNPs in 370 samples. Each color represents one of the six ancestral genomic components, and each vertical bar represents an individual. **C.** Genetic structure revealed by Bayesian Analysis of Population Structure (BAPS) based on the mitogenomes of 665 samples. Each color represents one of the seven BAPS groups, and each vertical bar represents an individual. **D.** PCA plot based on genomic SNPs of 370 samples, with PC1 plotted against PC2. **E.** Phylogenetic tree reconstructed by IQ-tree based on mitogenomic haplotypes. Table S23 presents individual information on haplotypes. Different colors represent seven different BAPS groups. Different regions are shown on the tree diagram. Red circles at the nodes represent bootstrap values larger than 70. **F.** Phylogenetic tree reconstructed by RAxML based on genomic SNPs in 370 samples. Different colors represent five different regions. All nodes have a bootstrap value of 100 except for these with grey circles.

PCA and phylogenetic analyses supported the general clustering of samples from different regions, except for the Japanese Kyushu populations, which clustered with the South Korean samples (Figures 3D, 3F, and S5). Three South Korean samples clustered with the Japanese Honshu and Chinese samples in the phylogenetic tree (Figure 3F). We also performed ADMIXTURE and PCA analyses separately for the five different regions to investigate the potential genetic structure within these populations (Figures S6 and S7). No admixture was detected in the American, South Korean, and European populations (K = 1). Two ancestry components were detected in Chinese and Japanese populations (K = 2, Figures S6A and S6B). A high degree of admixture was only detected in Chinese populations (Figures S6A and S7A), which was consistent with the results above on all populations (Figure 3B).

The lack of recombination and maternal characteristics resulted in a divergence in the genomic structure and phylogenetic relationships between the nuclear and mitochondrial datasets; however, the general patterns were similar. Bayesian Analysis of Population Structure (BAPS) resolved seven mitogenomic groups in the mitogenomic dataset (K = 7; Figure 3C; Table S12), corresponding to multiple lineages in two major clades of the phylogenetic tree (Figure 3E). In Clade A, Group 2 consisted of East Asian haplotypes, Group 3 of European haplotypes, and Groups 4 and 5 of Chinese and European haplotypes derived from Group 1, which included the American and Japanese Honshu haplotypes and was split into three lineages (Figures 3C and 3E). In Clade B, Group 7 comprised East Asian haplotypes derived from Group 6, including the American and Japanese Honshu haplotypes (Figures 3C and 3E).

TreeMix results based on genomic SNPs resolved two major clades diverged from American populations, containing the South Korean and Japanese populations, and European and Chinese populations, respectively (Figure 4E). Two migration events with high weight were identified, one from American to Japanese populations and another from South Korean to Chinese populations (Figure 4E). The residual plot indicated potential gene flow among American and European, and Japanese and South Korean populations (Figure 4F).

**Figure 4.**
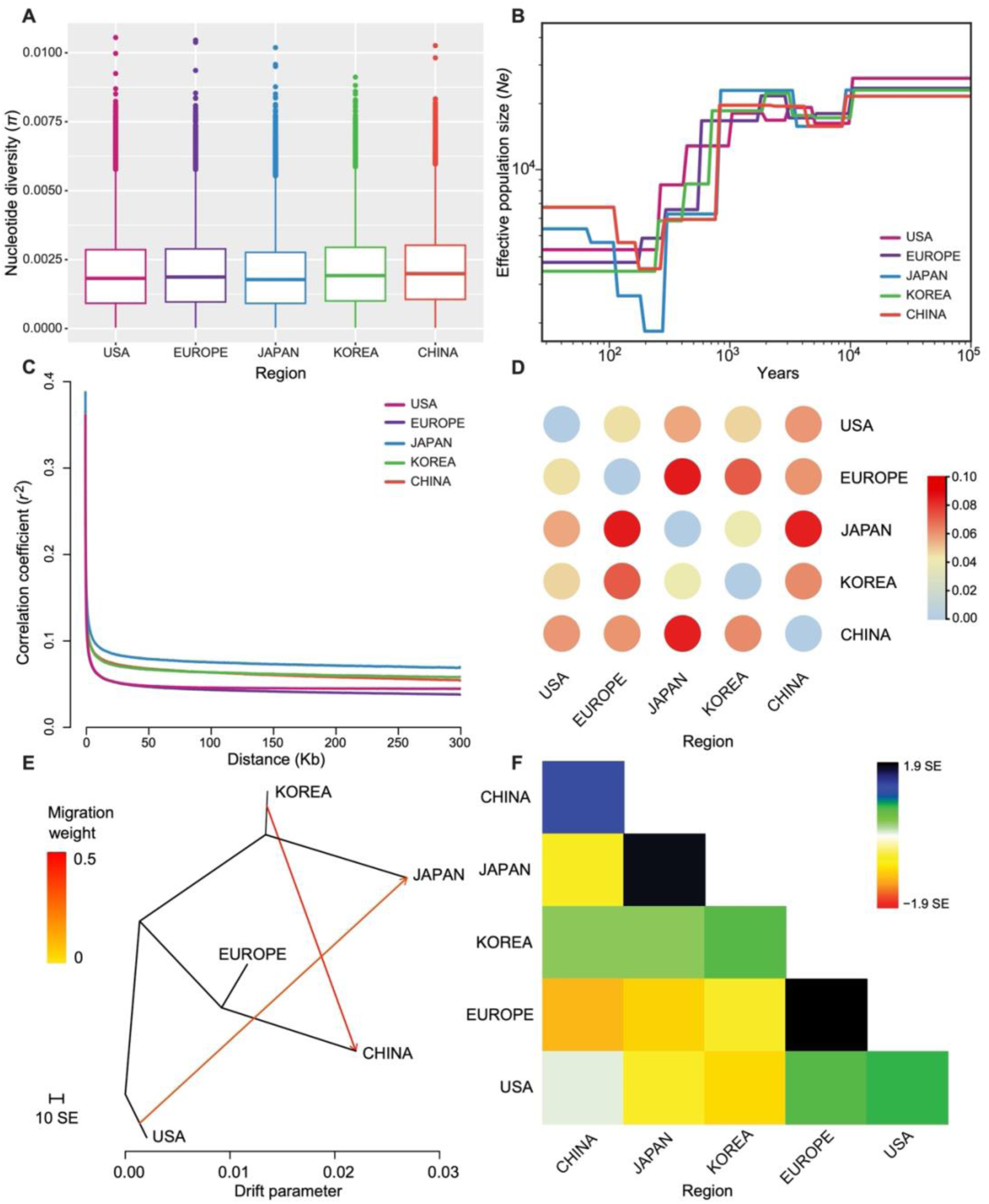
Genetic diversity, demographic history, and migration. **A.** Genome-wide mean nucleotide diversity under a 3-kb sliding window. **B.** Demographic history, depicted by the changes in effective population sizes (*Ne*) over time and inferred by SMC++. **C.** LD decay between SNPs as measured using the coefficient of correlation (*r^2^*) of alleles at any two loci. **D.** Heat map of Weir and Cockerham’s mean estimate of *F*_ST_ among five different countries and regions. **E.** TreeMix phylogeny with two migration events modelled. The horizontal branch lengths are proportional to the amount of genetic drift that has occurred along that branch. The scale bar shows 10 times the average S.E. of the entries in the sample covariance matrix. **F.** Residual fit of the observed versus the predicted squared allele frequency differences, expressed as the number of S.E. of the deviation. Residuals above zero represent sites that are more closely related to each other in the data than in the tree.

### Genetic diversity and demographic history

The genome-wide nucleotide diversity (π) calculated on sliding-windows was highest in the South Korean and Chinese populations (mean π = 0.0021), while the Japanese population had the lowest diversity (mean π = 0.0019; Figure 4A). When analyzing each sampling population based on all genomic SNPs, the highest observed heterozygosity (Ho) and π was found in Chongqing, China, while the lowest was in Tokushima, Japan (Table S13). South Korean populations had the highest diversity (mean Ho = 0.27, π = 0.27) and Japanese populations the lowest (mean Ho = 0.21, mean π = 0.21). Interestingly, when analyzing these populations based on only variant SNPs, the highest Ho and π was found in Georgia, USA, while the lowest Ho was in Qingdao, China and the lowest π was in Shanghai, China (Table S13). American population had the highest diversity (mean Ho = 0.36, mean π = 0.38), followed by the Japanese populations (mean Ho = 0.35, mean π = 0.36), while the Chinese and South Korean populations had the lowest (mean Ho = 0.33, mean π = 0.34; Table S13). For the mitogenomic dataset, the highest diversity was found in the South Korean (π = 0.00333) and American populations (π = 0.00323), while European population had the lowest (π = 0.00074; Table S1). These results indicated that Chinese and South Korean populations possess higher diversity on fixed SNPs, which supported that multiple introductions may have increased the genetic diversity in these populations.

Weir and Cockerham’s mean estimate of *F*_ST_ was generally lower for American populations (*F*_ST_ = 0.045–0.059) compared to other regions (*F*_ST_ = 0.060–0.085), except for South Korea and Japan (*F*_ST_ = 0.041), which had the lowest *F*_ST_. Chinese populations had relatively low *F*_ST_ with Europe (*F*_ST_ = 0.060) and South Korea (*F*_ST_ = 0.062) but high *F*_ST_ with Japan (*F*_ST_ = 0.084). The highest *F*_ST_ was observed between Europe and Japan (*F*_ST_ = 0.085; Figure 4D; Table S14). For the mitogenomic dataset, the highest pairwise *F*_ST_ was also found between Europe and Japan (*F*_ST_ = 0.729), followed by that between China and Japan (*F*_ST_ = 0.516), while the lowest *F*_ST_ was observed between the USA and South Korea (*F*_ST_ = 0.109), followed by that between the USA and China (*F*_ST_ = 0.135; Table S14).

We determined the demographic history of *C. ciliata* using both SMC++ and Stairway Plot 2. Our findings suggest that the effective population size (*Ne*) of *C. ciliata* began to rapidly decrease 1000 years ago, reaching its lowest level in the last 200–300 years (Figures 4B and S8). The Japanese population showed the longest LD decay distance, while the American and European population showed the shortest LD decay distance (Figure 4C).

### Genomic signatures of selection associated with invasion

To examine whether selection contributed to the successful invasion of *C. ciliata*, we scanned the genomes of all four invasive populations (Europe, Japan, South Korea, and China) as well as the native populations (USA). Given that the direction of selection was unknown, we applied both *F*_ST_ and θ_π_ ratios for detecting the genomic signatures of selection associated with invasion. We used a threshold of the top 1% for *F*_ST_ and preserved both the top and bottom 0.5% candidates for θ_π_ ratio. To avoid false-positive signatures and the effects of population structure, we only considered genomic loci shared by both methods as selection targets. As a result, we identified 7872 SNPs as genomic signatures of selection (Table S15), which were assigned to 237 genes (Table S16). Of these signatures, 18% (1420) of SNPs and 19.4% (46) of genes were shared between at least two comparisons of invasive and native populations (Figure 5E).

**Figure 5.**
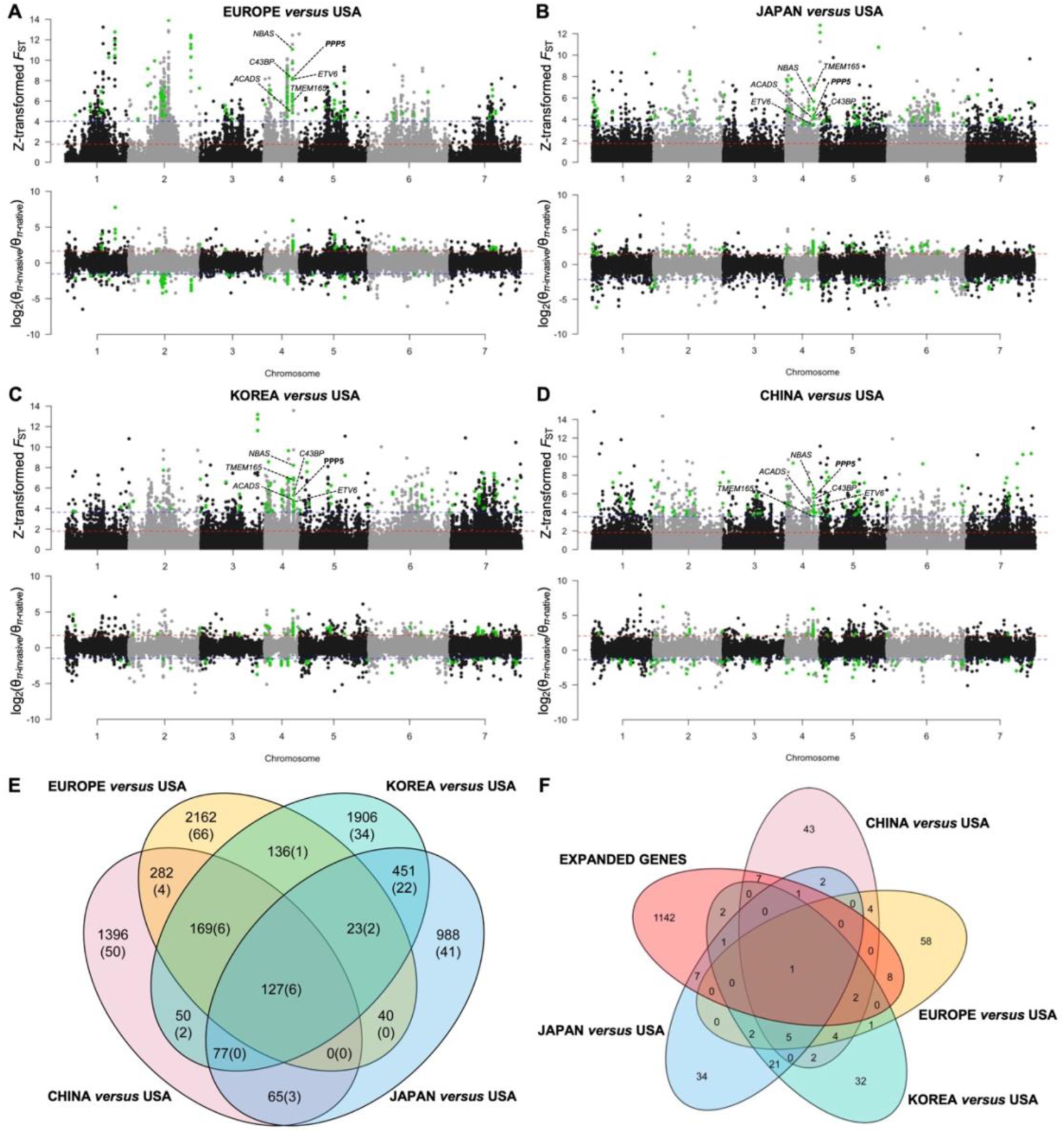
Genomic signatures of selection associated with invasion. **A–D.** The upper Manhattan plots show selection signatures detected by *F*_ST_. The x-axis represents seven chromosomes of the *C. ciliata* genome, and the y-axis represents the values of *Z*-transformed *F*_ST_. The horizontal dashed lines represent the thresholds of 1% (in blue) and 5% (in red) values. The bottom plots show selection signatures detected by nucleotide diversity, θ_π_ ratio (θ_π-invasive_/θ_π-native_). The y-axis represents log_2_(θ_π-invasive_/θ_π-native_). The horizontal dashed lines represent the thresholds of the top (in red) and bottom (in blue) 0.5% values. The highlighted dots represent the shared loci detected by two methods. **E.** Venn diagram showing overlapped SNPs (genes) between those identified in four pairs of comparison between invasive and native populations. **F.** Venn diagram showing overlapped genes between those identified in four pairs of comparison between invasive and native populations and expanded genes identified in comparative genomic analysis.

To investigate the potential effects of these signatures on the environmental adaptation of *C. ciliata*, we applied environmental association analysis (EAA) based on the 7872 SNPs and ten biological climatic variables which were selected after filtering out strongly corelated variables. With the criterion of log_10_ [Bayes factor (BF)] values > 1.5 [39,40], we found allele frequency variations in ∼4% SNPs (294) were correlated with environmental variation across 42 sampling populations (Table S17). Of these SNPs, 35.4% (104) correlated with maximum temperature of the warmest month (BIO5), followed by 15.0% (44) corelated with mean temperature of the wettest quarter (BIO8), and 11.6% (34) corelated with minimum temperature of the coldest month (BIO6), indicating their potential roles in temperature adaption of *C. ciliata*.

Notably, 127 shared SNPs on six genes were clustered on Chromosome 4 over a ∼47 kb genomic region. These genes include neuroblastoma-amplified sequence (*NBAS*), ETS variant transcription factor 6 (*ETV6*), transmembrane protein 165 (*TMEM165*), collagen type IV alpha-3-binding protein (*C43BP*), serine/threonine-protein phosphatase 5 (*PPP5*), and short-chain specific acyl-CoA dehydrogenase (*ACADS*; Figures 5A–E; Table S18). Higher nucleotide diversity was observed in this genomic region in invasive populations compared to native populations, indicating that they might be favored by diversifying selection (Figures 5A–D). To rule out bias owing to sample size, we performed the same analysis using 20 randomly selected samples from each region and obtained similar results (Table S19; Figure S9), except that one adjacent gene, chitooligosaccharidolytic β-N-acetylglucosaminidase (*BmChiNAG*), was included as a shared selection target in the cluster. Furthermore, *PPP5* was also included in the expanded gene set of the *C. ciliata* genome (Figure 5F).

### Widespread balancing selection signatures in native ranges

To determine the origin of adaptation, we investigated whether either the genomic signatures related to selection during an invasion or the expanded genes of this species were maintained in native populations by long-term balancing selection. We scanned the genomes of the native North American populations for signatures of long-term balancing selection using *β* scores and chromosomal SNPs with a MAF > 0.05 [19]. High *β* scores in this analysis indicated an excessive number of SNPs at similar frequencies.

Notably, we found that the top 1% of candidate SNPs with signatures of long-term balancing selection were widespread across the seven chromosomes (Figure 6A). We tested whether the genomic loci associated with invasion had higher signatures of long-term balancing selection relative to the whole genome in the native populations (Table S20). Across all seven chromosomes we found that the candidate SNPs had significantly higher *β* than the other genomic regions (mean *β* = 3.13 v. mean *β* = 1.58; Mann-Whitney U test, *U* = 3.8e9, *P* = 5.3e-51). When analyzing the chromosomes separately, we found candidate SNPs on Chromosome 2 (mean *β* = 4.56; Mann-Whitney U test, *U* = 1.9e8, *P* = 4.1e-84) and Chromosome 3 (mean *β* = 7.19; Mann-Whitney U test, *U* = 1.0e8, *P* = 6.4e-54) had significantly higher *β* than the other chromosomal regions (mean *β* = 1.61 for chromosome 2, mean *β* = 1.59 for Chromosome 3). Interestingly, we found 33 sites on Chromosome 2 and 40 sites on Chromosome 3 with the highest *β* scores are all identified as the candidate SNPs of selection associated with invasion (Figure 6A).

**Figure 6.**
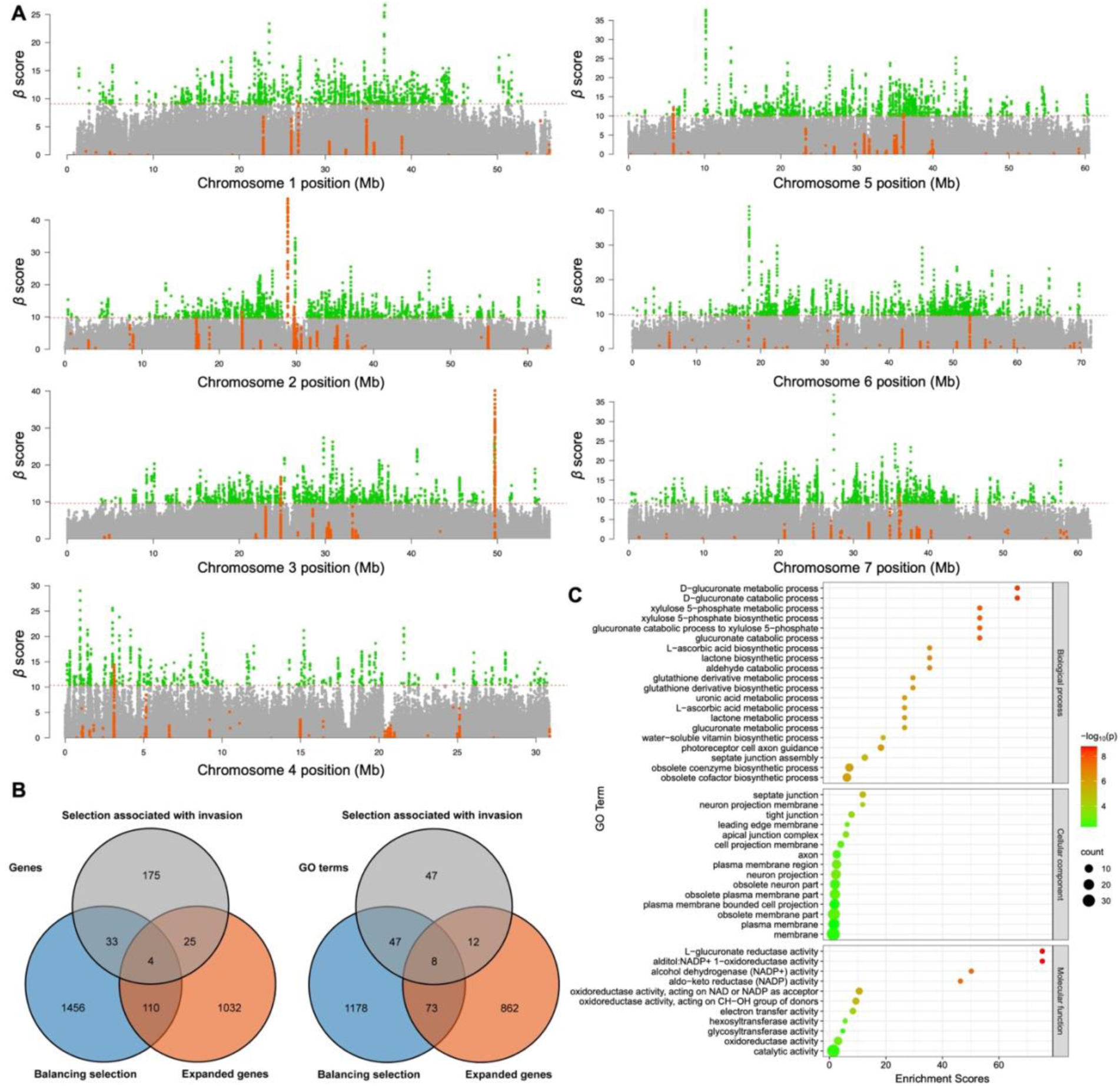
Genomic signatures of long-term balancing selection in native ranges. **A.** Manhattan plots showing balancing selection signatures in native populations detected by *β* scores on seven chromosomes of *C. ciliata* genome. The horizontal dashed lines represent the 1% thresholds of the *β* scores. Green dots show SNPs that have *β* scores above the 1% thresholds. Orange dots show candidate SNPs under selection and associated with invasion identified in four pairs of comparison between invasive and native populations. **B.** Venn diagrams showing overlapped genes and GO terms between those identified in balancing selection, selection signatures associated with invasion, and the expanded gene set. **C.** GO enrichment of shared genes between balancing selection and expanded gene set.

The top 1% of candidate SNPs with signatures of long-term balancing selection corresponded to 1603 genes (Table S21), which overlapped with both the selection signatures associated with invasion (15.6%) and the expanded gene set (9.7%; Figure 6B). The expanded gene set also overlapped with the selection signatures associated with invasion (12.2%; Figure 6B). To reflect the potential functions of these genes, we investigated the overlap between the significantly enriched GO terms in these gene sets. In total, 41.2% GO terms in the selection signatures associated with invasion and 8.5% in the expanded gene set were shared with balancing selection signatures (Figure 6B). Consistent with the GO enrichment of the expanded genes (Figure 2C), the genes shared with balancing selection signatures were enriched in oxidoreductase and metabolic activities (Figure 6C; Table S22). For instance, “L-glucuronate reductase activity” [GO:0047939], “alditol:NADP^+^ 1-oxidoreductase activity” [GO:0004032], “oxidoreductase activity, acting on the CH-OH group of donors, NAD or NADP as acceptor” [GO:0016616], “oxidoreductase activity, acting on CH-OH group of donors” [GO:0016614], “oxidoreductase activity” [GO:0016491], and “catalytic activity” [GO:0003824] were the top GO terms in the enrichment of expanded genes of the *C. ciliata* genome (Figures 2C and 6C).

## Discussion

Comprehensively studying the mechanisms underlying invasiveness is crucial to elucidate the evolutionary origins of adaptations that facilitate successful invasions. In the present study, we assembled a complete and highly continuous genome of *C. ciliata*, excluding the Y chromosome. This is the first assembly and chromosome-level reference genome of Tingidae. With ∼2,000 described species, lace bugs are distributed worldwide, are usually host-specific, and can be very destructive to plants. The novel reference genome assembled in this study filled a significant knowledge gap in the genomics of lace bugs and may provide a critical genomic reference for future research on this ecologically and economically significant group of insects.

### Gene family expansion and contraction in the *C. ciliata* genome

Based on comparative genomic analyses of ten hemipteran insects, we observed 276 expanded gene families associated with oxidoreductase and metabolism-related functions (Figures 2A and 2C; Tables S5 and S6). This is in contrast to observations made in many polyphagous insects whose invasiveness is attributed to an increased capacity to use novel nutritional sources. For instance, the gustatory receptor genes in the fall webworm [13] and the olfactory receptor genes in the codling moth [21] have expanded to facilitate their successful and rapid adaptation. *Corythucha ciliata* was unlikely to utilize novel nutrition sources or host plants to facilitate rapid and successful establishment; instead, the decisive factor in their distribution and invasion appears to be their capacity to adapt to a wide temperature range [41]. Previous physiological studies reported that high temperatures induced oxidative stress, leading to high anaerobic respiration and antioxidant defenses in *C. ciliata* in the laboratory and field [42]. This mechanism guards against oxidative damage caused by reactive oxygen species accumulation [42]. Strong enzymatic and metabolic activities, particularly the oxidoreductase activity, may provide high tolerance to oxidative stress under varying temperatures.

In the sycamore lace bug genome, we identified 228 contracted gene families (Figure 2A) significantly enriched for the ATP-binding cassette (ABC) transporter, mediation of the transport of a variety of physiologic lipid compounds (sterol, cholesterol, and other lipid), and for development and morphogenesis (Tables S7 and S8). The ABC transporter and cytochrome P450 gene families have been suggested to relate to the resistance of pesticide and xenobiotic in insects [21,43–46]; however, these two gene families exhibited opposite trends in their expansion and contraction during evolutionary history. The specific contraction of gene families in the *C. ciliata* genome might be related to the oligophagy of this lace bug and the long-term co-evolution with host sycamore trees, while particular resistance is needed and selected for, other resistances might be redundant and are lost during evolution.

### Global invasion history of *C. ciliata*

Based on a vast population genomic dataset with 2.74 million SNPs and a comprehensive sampling covering native and invasive ranges across three continents, we identified two major invasion routes of *C. ciliata*. During an invasion, a strong bottleneck and inbreeding may reduce the genetic diversity of founder populations [47–49]. The lowest genomic-wide nucleotide diversity was found in Japan and the lowest mitogenomic diversity in Europe; these findings are consistent with these effects (Figure 4A; Table S1). The largest genetic differentiation was observed between Europe and Japan (Figure 4D; Table S14), and the genetic structure and migration analyses (Figures 3B–F, 4E and 4F) supported that the European and Japanese Honshu populations were independently derived from two different invasion events. The Chinese and South Korean populations were derived from both invasion routes, indicating the existence of a contact zone in East Asia.

Interestingly, the Chinese and South Korean populations exhibited higher nucleotide diversity than both the native American populations and earlier introduced European populations [25,26] (Figure 4A; Table S1), although the native American population possessed the highest genetic diversity of variant sites (Table S13). This was possibly caused by genetic admixture of different invasive sources; nevertheless, the larger LD decay distance found in the Chinese and South Korean populations was consistent with their more recent invasion history (Figure 4C). Multiple introductions have been reported to ameliorate the loss of genetic diversity during invasion and may facilitate the fitness of invasive populations [49–51].

Understanding the impact of anthropological activities on biological invasion and population dynamics can inform strategies for managing invasive species and conserving biodiversity. Given the limited flight capacity of *C. ciliata*, long-distance dispersion, particularly across continents, was possibly accomplished through anthropogenic activities. Urban afforestation facilitated international and domestic transportation of sycamore trees, providing immense opportunities for this insect to spread across geographic barriers [31]. East Asia, particularly regions of eastern China, South Korea, and Japan, has been identified as a hotbed of invasion in the present and previous studies [37]. Further, we observed a drastic population contraction from 1000 to 200 years ago, which may have been linked to the exploitation and urbanization of North America, along with the possible logging of sycamore trees, a key source of wood at the time.

### Genomic signatures of selection associated with invasion

Environmental differences between native and invasive ranges might result in divergent selective pressures and genomic changes of critical loci in novel habitats may enhance the fitness and invasiveness of invasive populations [12–14]. In this study, we examined the hypothesis that selection may enhance the thermal adaptation of populations in invasive ranges and lead to varying success compared to native populations. Previous studies reported that invasive species populations are sometimes more successful than native populations due to a higher thermal tolerance as global climate change drives temperatures rise [3]. Based on the selection signature and environmental association analyses conducted in our study, we found a number of candidate SNPs (∼4%) under selection and associated with invasion were corelated with environmental variables, especially the highest and lowest temperature across their distribution ranges (Table S17). These results supported our hypothesis that selection may enhance the temperature adaptation of invasive populations, contributing to their higher thermal tolerance and rapid invasion.

Notably, *PPP5* and adjacent gene blocks were repeatedly selected during multiple invasion scenarios (Figures 5A–D). *PPP5* is involved in various cellular signaling pathways in higher eukaryotes, including those initiated by atrial natriuretic peptides, oxidative stress, and glucocorticoids, and pathways operating through G-proteins [52,53]. Additionally, through its regulatory tetratricopeptide repeat domain, *PPP5* interacts with and binds to both *HSP70* and *HSP90*, which have been found to enhance the stress tolerance of *C. ciliata* and to contribute to the invasive success of many other species [3,53–55]. *PPP5* and *ETV6*, both found within this gene block, are involved in the mitogen-activated protein kinase (MAPK) signaling pathway in the KEGG database. MAPK signaling cascades use protein phosphorylation to convey signals intracellularly and contribute crucially to oxidative stress and metabolism [56]. This close genetic linkage might be favored by selection with reduced recombination [20,57–59]. This gene block was repeatedly selected during independent invasions of *C. ciliata* and might play a key role in increasing their thermal tolerance in invasive ranges, resulting in varying success. This may also have facilitated the extensive invasion of numerous subtropical regions by this temperate species, despite the rise in temperature rise attributed to climate change [7,31].

### Widespread balancing selection signatures in native ranges

Fluctuating environments may induce balancing selection, favoring multiple alleles at a given locus to retain functional polymorphisms [12,19]. *Corythucha ciliata* span a wide latitudinal range and have multiple overlapping generations yearly. These bugs overwinter as adults under crevices, leaf litter, and loose bark [25,60]. Thus, *C. ciliata* frequently experience temperature fluctuation, including hot summers (36°C to 40 °C) and cold overwintering (–30°C to –10 °C) [61–63]. As far as we understand, balancing selection is the major force that maintains multiple alleles at the loci of a population when one allele is fitter under one condition and another allele is fitter under another condition. Given the seasonally fluctuating environments *C. ciliata* is facing, we believe that signatures of long-term balancing selection are likely to be detected in native populations [64,65]. In accordance with this theory, the highest genetic diversity of variant SNPs was found in the native American populations of *C. ciliata* (Table S13). Balancing selection is common in nature; however, research on its role in maintaining the crucial standing variation that natural selection could act on, particularly in populations experiencing rapid environmental changes during invasion, is scarce.

In this study, we were particularly interested in the long-term balancing selection acting on the whole genome in the native ranges [19], although the balancing selection on a few specific loci could also allow a species to overcome bottleneck during invasion [66]. We found widespread balancing selection signatures along the seven chromosomes (Figure 6A) in the native ranges. Moreover, candidate SNPs under selection and associated with invasion showed significantly higher signatures of long-term balancing selection compared with other genomic regions (Table S20, Figure 6A). This further supported our hypothesis that these loci under balancing selection in native ranges may play pivotal roles in the establishment of *C. ciliata* after introduction into the invasive ranges. The genes under long-term balancing selection overlapped substantially with both expanded genes and genes under selection and associated with invasion (Figure 6B). These genes exhibited specific enrichment in oxidoreductase and metabolism-related functions (Figures 4 and 6B; Tables S7 and S8), despite the highly polygenic mechanism of genetic redundancy-mediated thermal adaptation [67]. This adds further support to the concept that the pre-introduction adaptation of this species originated from long-term balancing selection in its native ranges, potentially contributing to its oxidative stress and thermal tolerance and capacity to invade and establish novel habitats.

## Conclusions

In this study, we assembled the first Tingidae genome, presenting a chromosome-level genome of the sycamore lace bug *C. ciliata*. High-coverage PacBio, Illumina, and Hi-C sequencing were employed. We explored species-specific gene family expansions and contractions as well as in-depth resequencing of 402 samples covering both native and invasive ranges across three continents. We identified two independent invasion routes of this lace bug, both of which originated from the USA and subsequently spread through either Europe or Japan, forming a contact zone in East Asia. We identified genomic signatures of selection associated with invasion and long-term balancing selection in the native ranges. These genomic signatures overlapped with each other and the expanded genes in the *C. ciliata* genome, which were associated with oxidoreductase and metabolic activities. This is suggestive of improved oxidative stress and thermal tolerance in this lace bug. Our study sheds light on the genomic architecture of extraordinary invasive capacity of *C. cliliata*, providing insights into the mechanisms underlying the successful expansion of invasive species in rapidly changing environments.

It should be noted that several genes and loci were identified as potential targets of selection; however, direct validation of their functions is complicated by the lack of gene silencing and knockout systems in *C. ciliata* and the complexity of the such experiments in this small host-specific herbivore. Nevertheless, our study may benefit developmental biology research as it presents the first high-quality Tingidae genome resource. Future studies may provide a more optimized solution for validating the genomic mechanism of adaptation in this species.

## Materials and methods

### DNA extraction and whole genome sequencing

*Corythucha ciliata* were obtained from China Agricultural University, Western Campus (40.0°N, 116.3°E). Given that *C. ciliata* possesses an XX/XY sex-determination system [34,35], only female samples were used for genome sequencing to lower the effects of heterozygosity and sequencing coverage on the quality of genome assembly. DNA extraction was performed with a DNeasy Blood & Tissue Kit (catalog No. 69504, QIAGEN, Hilden, Germany), and 26.7 μg of HMW DNA was obtained from 57 female samples. For short-read sequencing, a 350-bp insert size library was constructed and sequenced on the NovaSeq 6000 platform (Illumina, San Diego, CA) in 150 paired-end (PE) modes. For continuous long-read (CLR) sequencing, the 20-kb insert size SMRTbell library was constructed and sequenced on the Sequel II platform (PacBio, Menlo Park, CA) at Berry Genomics (Beijing).

The Hi-C sequencing library was subsequently prepared. For crosslinking, 108 female samples were fixed with 2% formaldehyde, and the crosslinked DNA was digested using *MboI* restriction endonuclease. A standard Illumina library preparation protocol was performed with the ligated DNA sheared to 300–600 bp. The Hi-C library was also sequenced on the NovaSeq 6000 platform in 150 bp PE modes.

For full-length transcriptome sequencing, 130 female samples were used. TRNzol Universal Reagent (catalog No. 4992730, TIANGEN, Beijing) was applied to extract total RNA, and 3.42 μg RNA was obtained, followed by constructing a 1–10 kb + 4–10 kb SMRTbell library. Circular consensus sequencing (CCS) was performed on the Sequel II platform. Raw sequencing data were filtered using the standard IsoSeq3 protocol (https://github.com/ylipacbio/IsoSeq3).

### Chromosome-level genome assembly

Illumina sequencing reads were used to estimate the genome size. JELLYFISH [68] was used to analyze the distribution of k-mer. GenomeScope v1.0 [69] was used to estimate genome size, heterozygosity, and repetitive sequences based on a k-mer distribution (k = 19).

For genomic assembly, the long PacBio CLR were corrected and trimmed using Canu v1.8 [70]. Wtdbg2 [71] was then used to assemble the trimmed reads (genome size = 400 Mb). To polish the initial assembly with PacBio long reads and Illumina short reads, Arrow (https://github.com/PacificBiosciences) and Pilon (https://github.com/broadinstitute/pilon), respectively, were used. Redundans v0.13c [72] was used to remove the duplicates, producing an improved assembly.

For chromosome-level scaffolding, Juicer [73] and 3D-DNA [74] were used to anchor the initial assembly onto seven chromosomes with the Hi-C reads. The sex chromosome was not specifically identified. BUSCO v3.0.1 was used to assess the completeness of the reference genome based on genes in the insecta odb10 database [75].

### Genome annotation

RepeatMasker v4.07 (http://www.repeatmasker.org) was applied to search for the repetitive sequences and transposable elements in the *C. ciliata* genome depending on a search in Repbase, and a *de novo* repeat library constructed using RepeatModeler v1.0.11 (http://www.repeatmasker.org/RepeatModeler). RepeatProteinMask and TRF were used to search for TE proteins and tandem repeats.

Protein coding gene structures were predicted using homology-, *ab initio*-, and transcriptome-based predictions. For homology-based predictions, GeMoMa v1.6 [76] was applied to the protein sequences extracted from the published genomes of *Halyomorpha halys*, *Diuraphis noxia*, *Oncopeltus fasciatus*, and *Rhodnius prolixus*. For *ab initio* prediction, Augustus v3.3 [77], SNAP [78], GlimmerHMM v3.0.4 [79], and GeneMark-ET v4.21 [80] were used. For transcriptome-based prediction, the full-length transcripts obtained were combined with reassembled transcripts based on available short-read RNA-seq data, which were mapped onto our reference genome using HISAT v2.0.4 [81] and Cufflinks v2.2.1 [82]. The intact open reading frames (ORF) were predicted using PASA v2.0.1 [83]. Ultimately, we integrated the annotations from these three strategies using EvidenceModeler (EVM) v1.1.1 [84].

Functional annotation of protein-coding genes was based on homolog searches and the most optimal matches to five public databases. GO, KEGG, and eggNOG were annotated using the eggNOG-mapper v2.1.9 [85]. Swiss-Prot and NCBI NR annotations were performed using BLASTP (e-value < 1e-5).

MCScan in JCVI [86] was applied to identify collinear blocks within the *C. ciliata* genome using the protein-coding gene sequences.

### Comparative genomic analyses

OrthoFinder v2.5.4 [87] was used to identify orthologous gene families. The protein sequences of 10 other paraneopteran species were downloaded from the GenBank database. *Frankliniella occidentalis* were used as the outgroup because Thysanoptera (thrips) is strongly supported as sister to Hemiptera based on phylogenomic analysis [88]. First, alternative splicing for each gene was filtered out, maintaining the longest transcript. We aligned the proteins between *C. ciliata* and 10 other paraneopteran species by BLASTP (e-value < 1e-5).

MAFFT v7.313 and its L-INS-i algorithm [89] were used to align the protein sequences of the identified single-copy genes. RAxML v8.0.19 [90] was used to construct the phylogenetic tree with 100 bootstrap replicates. MCMCTree in PAML v4.9 [91] was used to estimate divergence time with a two-node calibration (confidence interval) obtained from the TIMETREE database (https://timetree.org). CAFÉ5 [92] was applied to compare the generated gene family clusters. TBtools v1.112 [93] was used to perform GO and KEGG pathway enrichments for gene families exhibiting expansion and contraction in the *C. ciliata* genome.

### Population sampling, resequencing, SNPs calling, and mitogenomic assembly

In total, 410 *C. ciliata* specimens were collected from 42 populations (n = 42) in 10 countries on three continents: China (n = 19), South Korea (n = 5), and Japan (n = 4) in East Asia; the United States (n = 4) in North America; and Italy (n = 2), Germany (n = 2), Switzerland (n = 2), France (n = 2), Austria (n = 1), and Belgium (n = 1) in Europe. These populations cover both the invasive and native regions of the sycamore lace bug. All samples were collected from the sycamore leaves and stored in absolute ethanol in a −80 °C freezer. Another 259 specimens from 35 of the 42 populations and an additional set of four novel populations in China, which were not covered above, were used only for mitogenomic assembly. The 259 individuals were pooled with equimolar-quantity genomic DNA from two other species [37]. Genomic DNA was isolated using a DNeasy Blood and Tissue Kit, used to construct 350-bp insert size libraries, and were sequenced on the Illumina NovaSeq 6000 platform in 150 PE mode to obtain 8 Gb (∼20×) sequencing reads.

BWA-MEM v0.7.12-r1039 (https://github.com/lh3/bwa) was used to map the clean reads onto the assembled reference genome. The genome mapping results of each sample were sorted and converted into the BAM format using SAMtools v1.16.1 [94]. Duplicate reads were removed using Picard v2.21.6 (https://github.com/broadinstitute/picard). SNPs were called using GATK v4.1.9.0 [95], and all GVCFs were merged for joint genotyping to generate raw SNPs. VCFtools v0.1.15 [96] was used to filter raw SNPs. Only high-quality biallelic SNPs were retained for subsequent analysis, with missing rates < 0.15, minor allele frequencies > 0.05, and mean depths > 10.

Clean reads were also mapped onto the published reference mitogenome using Geneious Prime (https://www.geneious.com) to generate mitogenomes of all samples, with a ≤ 2% mismatch, ≤ 5-bp gap, ≥ 40-bp overlap, and ≥ 95% similarity.

### Population genetic structure

For the genomic SNPs, KING v2.2.5 [97] was first used to calculate the kinship coefficient between individual pairs and exclude samples that were predicted to be full-siblings of other samples within the same population (under a threshold of 0.177). To avoid the LD effects on the inference of genetic structure, genomic SNPs were pruned, and a single SNP was maintained per 3-kb window to generate independent loci using PLINK v1.90b6.18 [98]. SNPs associated with the sex chromosome were not excluded in our analyses. ADMIXTURE v1.3.0 [99] was used to infer the population genetic structure with independent SNPs. Hypothetical ancestral cluster K, with K ranging from 2 to 10, were analyzed to determine the optimum number. Principal component analysis (PCA) was performed using the PLINK software. RAxML was used to reconstruct the phylogenetic relationships of all samples based on the maximum-likelihood (ML) method with the GTR + G model. TreeMix 1.13 [100] was used to infer patterns of historical splits and admixture events among different regions through reconstructing the bifurcating ML tree with 100 bootstraps. PLINK was used to generate allele frequency data. The migration edges (m = 2) were modelled from 0 until when 99.8% of variance of relatedness between populations can be explained by the model (Figure S10).

For the mitogenomes, protein-coding gens, tRNA genes, rRNA genes and the control region were aligned separately using MAFFT. Bayesian analysis of the population structure was performed using Bayesian Analysis of Population Structure (BAPS) v6.0 [101] under the spatial clustering of groups of individuals. Phylogenetic analysis was performed based on the haplotypes generated by DnaSP v6.0 [102] (haplotype information shown in Table S23). We performed ML analysis using the ultrafast bootstrap approximation approach in IQ-TREE v1.6.5 [103] with 1,000 replicates.

### Genetic diversity and demographic history

VCFtools was used to calculate the genome-wide nucleotide diversity (θ_π_) of populations from five different regions, and Weir and Cockerham’s mean estimate of *F*_ST_ based on the sliding-windows along the genome were calculated. The observed homozygosity, observed heterozygosity, expected homozygosity, expected heterozygosity, and genetic diversity index (π) were calculated for each sampling population using the populations function in the Stacks 2.65 [104]. Mitogenomic diversity was calculated using DnaSP. Arlequin 3.5 [105] was used to calculate pairwise *F*_ST_ values to measure mitogenomic differentiation between different regions.

Twenty randomly selected samples representing every population in the five native and invasive regions (Table S19) were used to estimate the changes in effective population size (*Ne*) in the past using SMC++ [106]. The Stairway plot 2 method was suggested to be more robust when reconstructing more recent demographic histories, thus Stairway plot 2 [107] was also used to infer the demographic history. EasySFS was used to convert VCF format to the site frequency spectrum (https://github.com/isaacovercast/easySFS#easysfs). We applied the mutation rate of *Drosophila melanogaster*, 2.8 × 10^-9^ per base pair per generation, and a generation time of 0.2 because five generations per year can be completed in the wild for *C. ciliata* [30]. We calculated the correlation coefficient (*r^2^*) following increasing decay distance and plotted LD decay curves using PopLDdecay [108] to measure the degree of LD among populations in each region based on the biallelic SNPs filtered out without missing data.

### Selection signature detection

Genomic signatures of selection associated with invasion were analyzed based on filtered nucleotide SNPs using genome-wide distribution of population fixation statistics *F*_ST_ and nucleotide diversity θ_π_. We selected four pairs of invasive and native populations and used VCFtools to calculate *F*_ST_ and θ_π_ with 10 kb sliding window and 5 kb step sizes. Negative *F*_ST_ values were considered to be 0. For the results of θ_π_, we further calculated log_2_ (θ_π-invasive_/θ_π-native_). The divergence regions from the intersection of the top 1% *Z*-transformed *F*_ST_ and log_2_ (θ_π- invasive_/θ_π-native_) were regarded as the putatively selection signatures. To ascertain whether the number of samples biased the results, the same analyses were performed using 20 randomly selected samples from each region (Table S19).

We scanned balancing selection signatures in the native populations to test whether SNPs and genes that showed signatures of selection in the invasive populations or critical genes expanded in the *C. ciliata* genome were also maintained by balancing selection within native ranges. SNPs (MAF < 0.15) were filtered out to prevent false positives. Seven chromosomes were analyzed separately using BetaScan [109] to obtain *β* scores along the genome. Statistical analyses were conducted in R to compare the *β* scores of candidate SNPs with the other genomic regions.

SnpEff v4.3t [110] was used to annotate the genomic SNPs and selected regions based on the annotation of the *C. ciliata* reference genome.

### Environmental association analysis

Nineteen biological climatic variables were used in our environmental association analysis (EAA) to investigate the potential effect of candidate SNPs on the environmental adaptation of *C. ciliata*. We used raster 3.5.2 and SpatialPoints 1.4.6 packages in R to retrieve the corresponding climatic data for 42 sampling localities (Table S1) from the WorldClim database.

To avoid the multi-collinearity of highly corelated variables, 19 variables were first filtered based on Pearson’s correlation tests using IBM SPSS Statistics 19.0 (Chicago, IL, USA). The independent variables that were highly correlated (|r|≥0.85) were removed, and the following variables were used for this analysis: BIO1, annual mean temperature; BIO2, mean diurnal range; BIO3, isothermality; BIO4, temperature seasonality; BIO5, maximum temperature of the warmest month; BIO6, minimum temperature of the coldest month; BIO8, mean temperature of the wettest quarter; BIO12, annual precipitation; BIO13, precipitation of the wettest month; and BIO14, precipitation of the driest month. The environmental variables of populations were calculated as the absolute difference between the value of each variable and the average across all populations, standardized by the standard deviation. Bayenv2 [111] was used to investigate the correlation between environmental variables with standardized allele frequencies among populations due to its lower false positive rate and decent performance handling hierarchical population structure [112].

## CRediT author statement

**Zhenyong Du:** Methodology, Formal analysis, Investigation, Resources, Visualization, Writing - Original Draft, Writing - Review & Editing. **Xuan Wang:** Formal analysis, Investigation. **Yuange Duan:** Formal analysis, Investigation. **Shanlin Liu:** Investigation, Writing - Review & Editing. **Li Tian:** Methodology, Writing - Review & Editing. **Fan Song:** Resources, Validation. **Wanzhi Cai:** Conceptualization, Supervision, Writing - Review & Editing. **Hu Li:** Conceptualization, Project administration, Funding acquisition, Writing - Review & Editing.

## Competing interests

The authors have declared no competing interests.

## Acknowledgments

This work was supported by the National Natural Science Foundation of China [grant numbers 32170453 and 31922012]; the International Postdoctoral Exchange Fellowship Program from the China Postdoctoral Council [grant number PC2021094]; and the 2115 Talent Development Program of China Agricultural University. The authors wish to thank Renfu Shao, Xingyue Liu, Yuzhou Du, Liangming Cao, Haoyu Liu, Mingyi Tian, Guoquan Wang, Shuangmei Ding, Haoyong Ouyang, Yingqi Liu, Han Xu, Yan Li, Zhiteng Xu, Yuanyuan Wu, Zhuo Chen, Yunfei Wu, Zhiping Wang, Shixiang Jiang, Ruojin Tian, Yunchuan He, Kazutaka Yamada, Jun Soma, and many other colleagues and citizen scientists for their help in sample collection.

