## Supplementary material for "Global invasion history and genomic signatures of adaptation of a highly invasive lace bug": Figure S1-S10

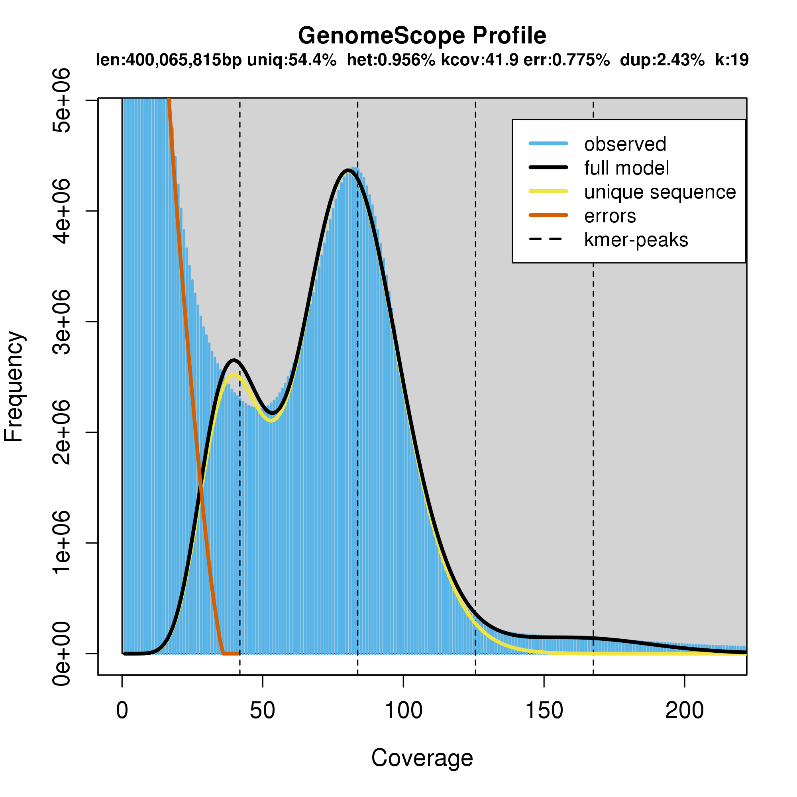

**Figure S1. K-mer coverage frequency plot for the *Corythucha ciliata*.** Analysis reveals an estimated genome size of 400 Mb, with a heterozygosity rate of 0.96% and 45.6% repetitive sequences.

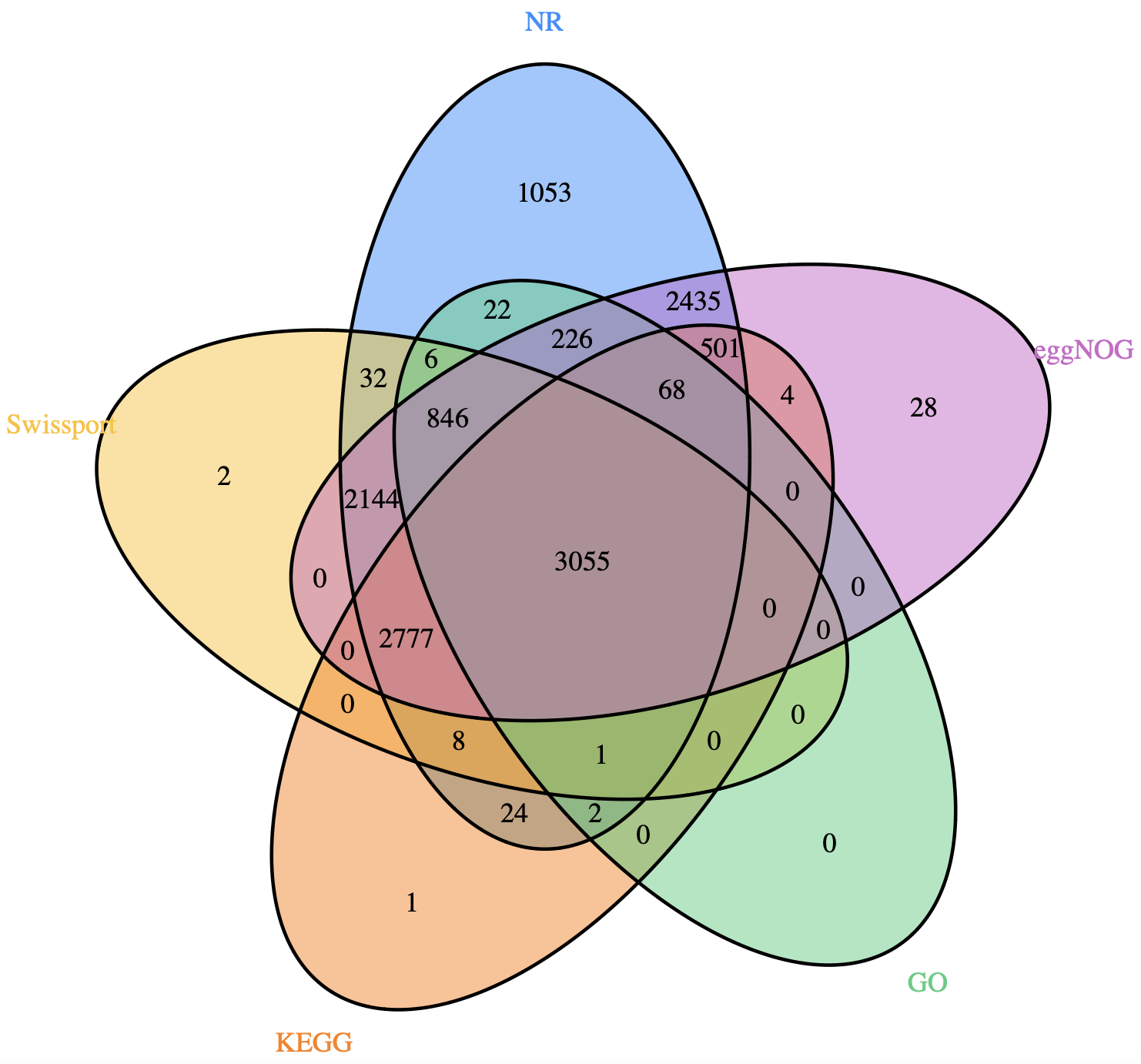

**Figure S2. Integrated functional annotation of the *Corythucha ciliata* genome using five databases.** The analysis shows that 13,235 genes (86.63%) were annotated by at least one database, with specific annotations as follows: NR database (13,200 genes, 86.40%), SwissProt (8,871 genes, 58.06%), KEGG (6,441 genes, 42.16%), GO (4,226 genes, 27.66%), and eggNOG (12,084 genes, 79.09%).

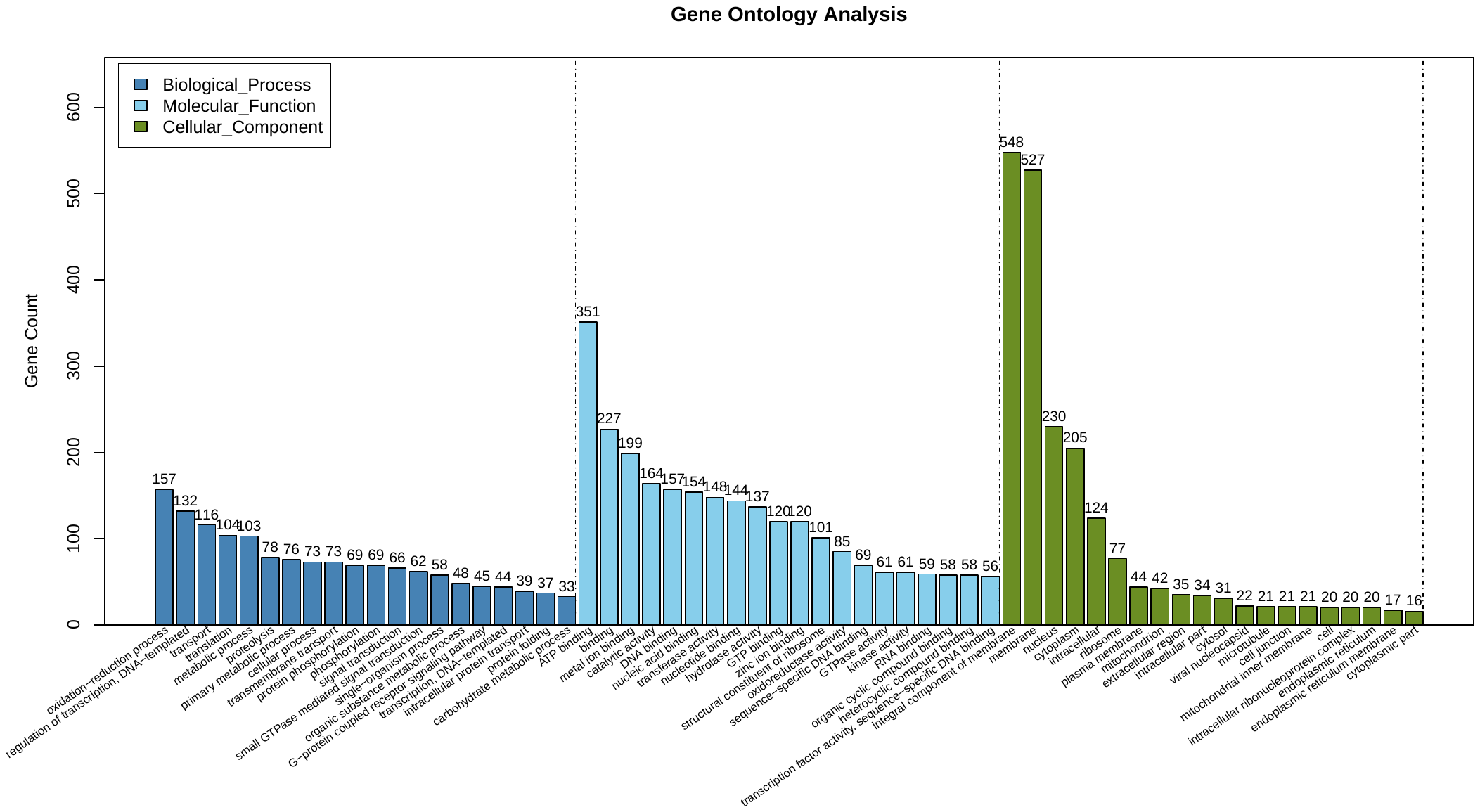

**Figure S3. Gene Ontology (GO) annotation of protein-coding genes in the *Corythucha ciliata* genome.**

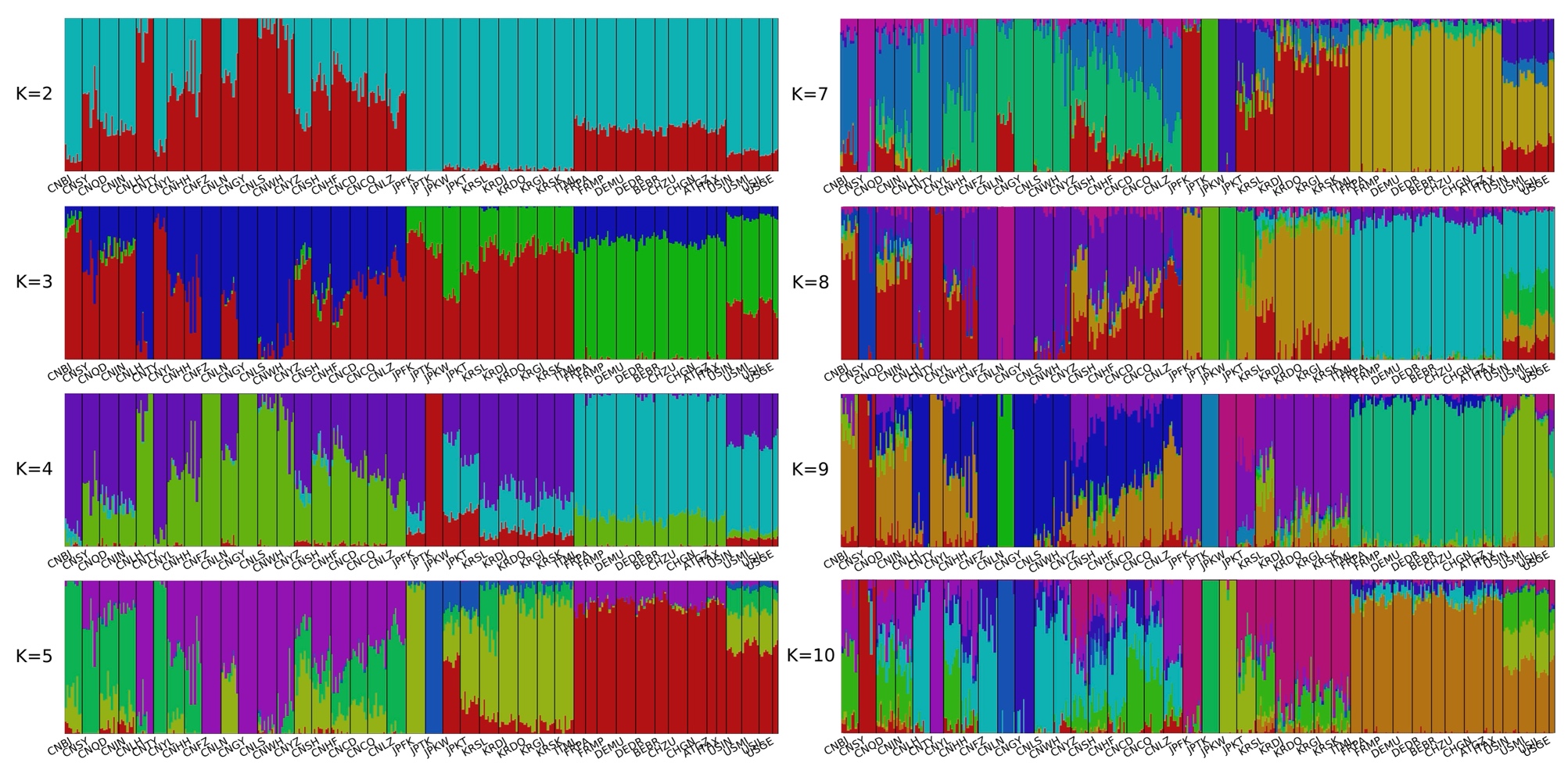

**Figure S4. Genetic structure of 370 *Corythucha ciliata* samples as revealed by ADMIXTURE analysis using genomic SNPs, for K = 2–10.** The corresponding Cross-Validation (CV) errors for each K value are: CV error for K = 2 is 0.54354, K = 3 is 0.52886, K = 4 is 0.51886, K = 5 is 0.51416, K = 6 (the lowest) is 0.51208*, K = 7 is 0.51251, K = 8 is 0.51258, K = 9 is 0.51293, and K = 10 is 0.51229. The lowest CV error at K = 6 suggests it as the most suitable model for the genetic clustering of these samples.

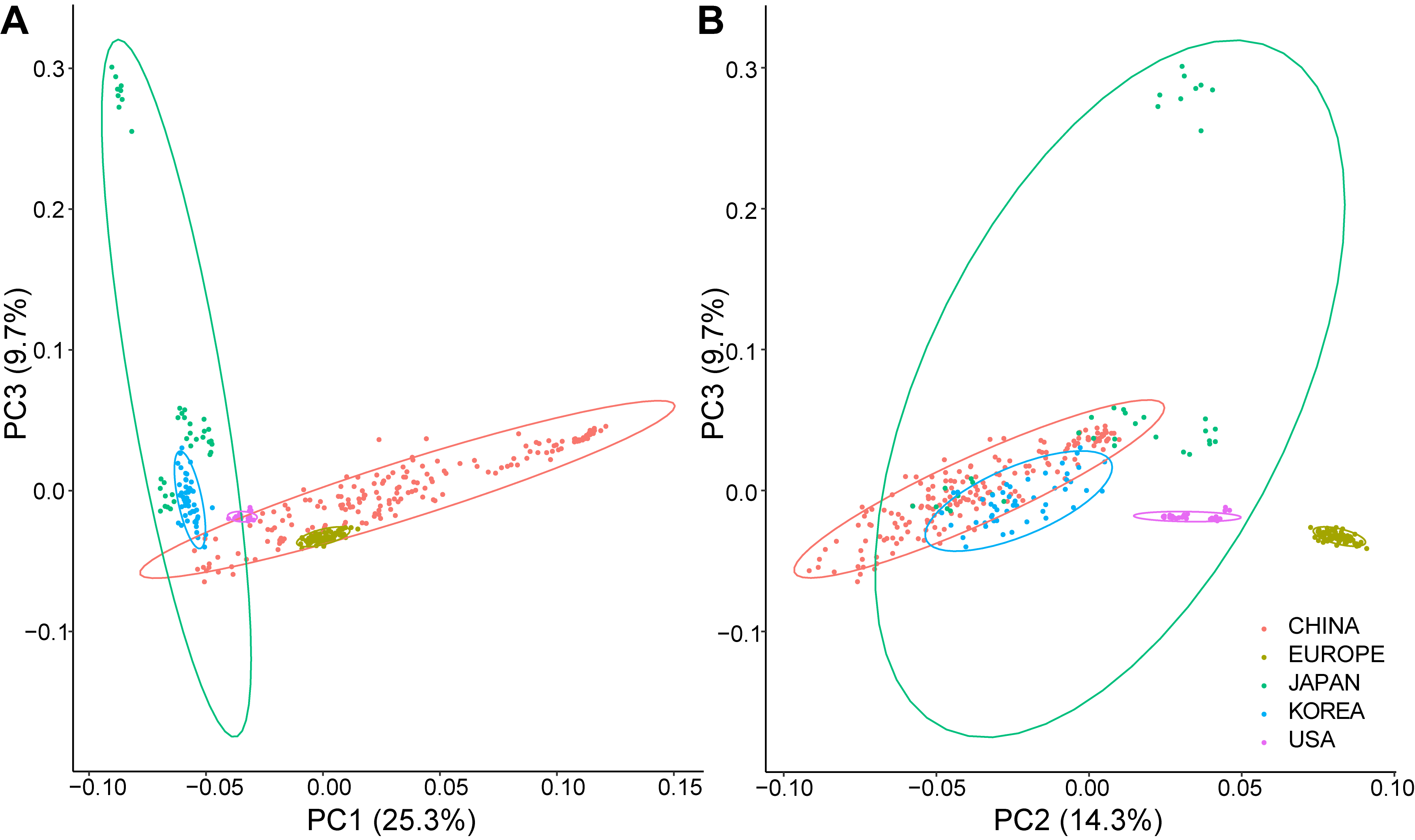

**Figure S5 Principal component analysis (PCA) plots based on genomic single nucleotide polymorphisms (SNPs) of 370 samples** Panel A shows PC1 plotted against PC3, while Panel B shows PC2 plotted against PC3. Sample origins are abbreviated as follows: CN for China, EU for Europe, JP for Japan, KR for South Korea, and US for the United States.

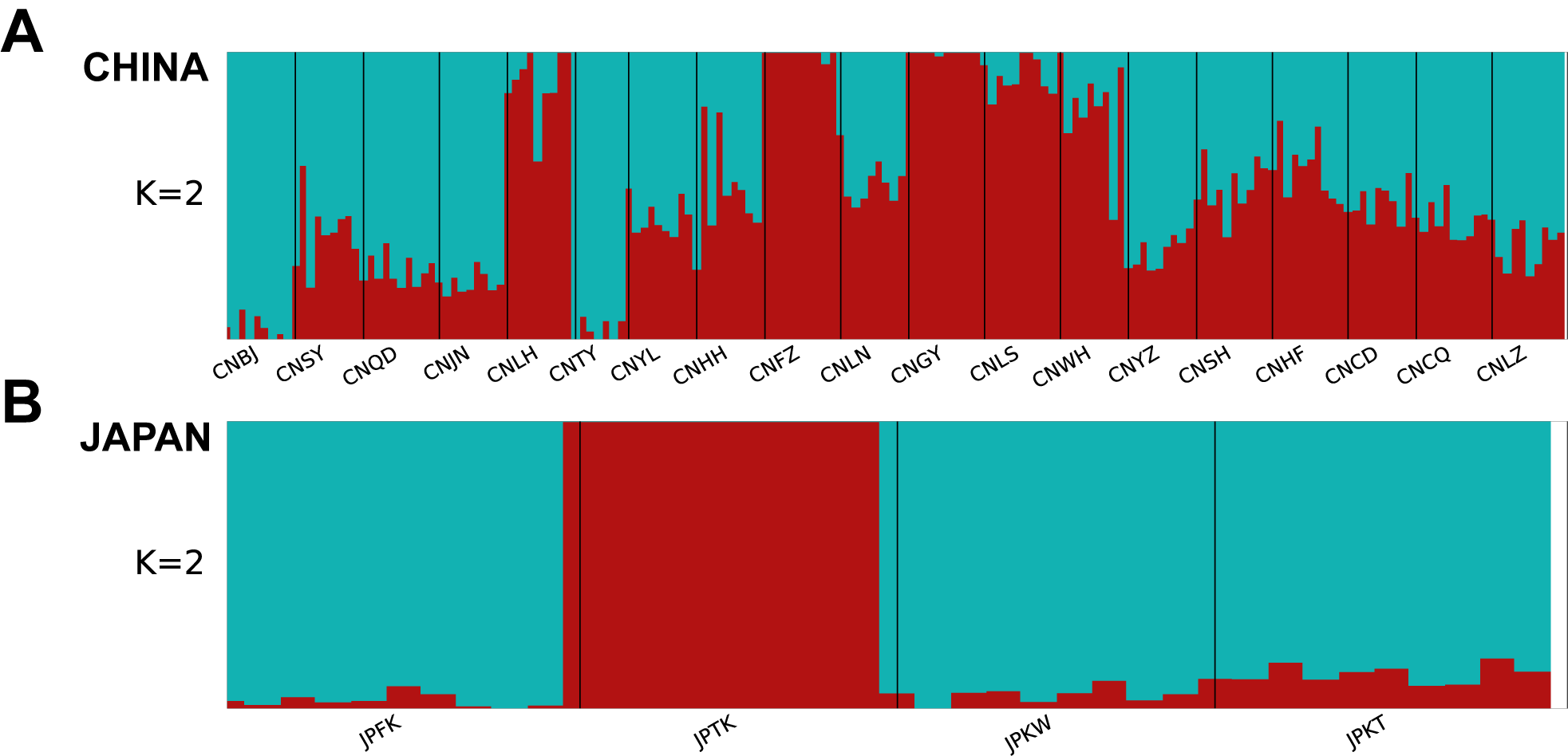

**Figure S6 Genetic structure of Chinese and Japanese populations as revealed by ADMIXTURE analysis using genomic SNPs.**

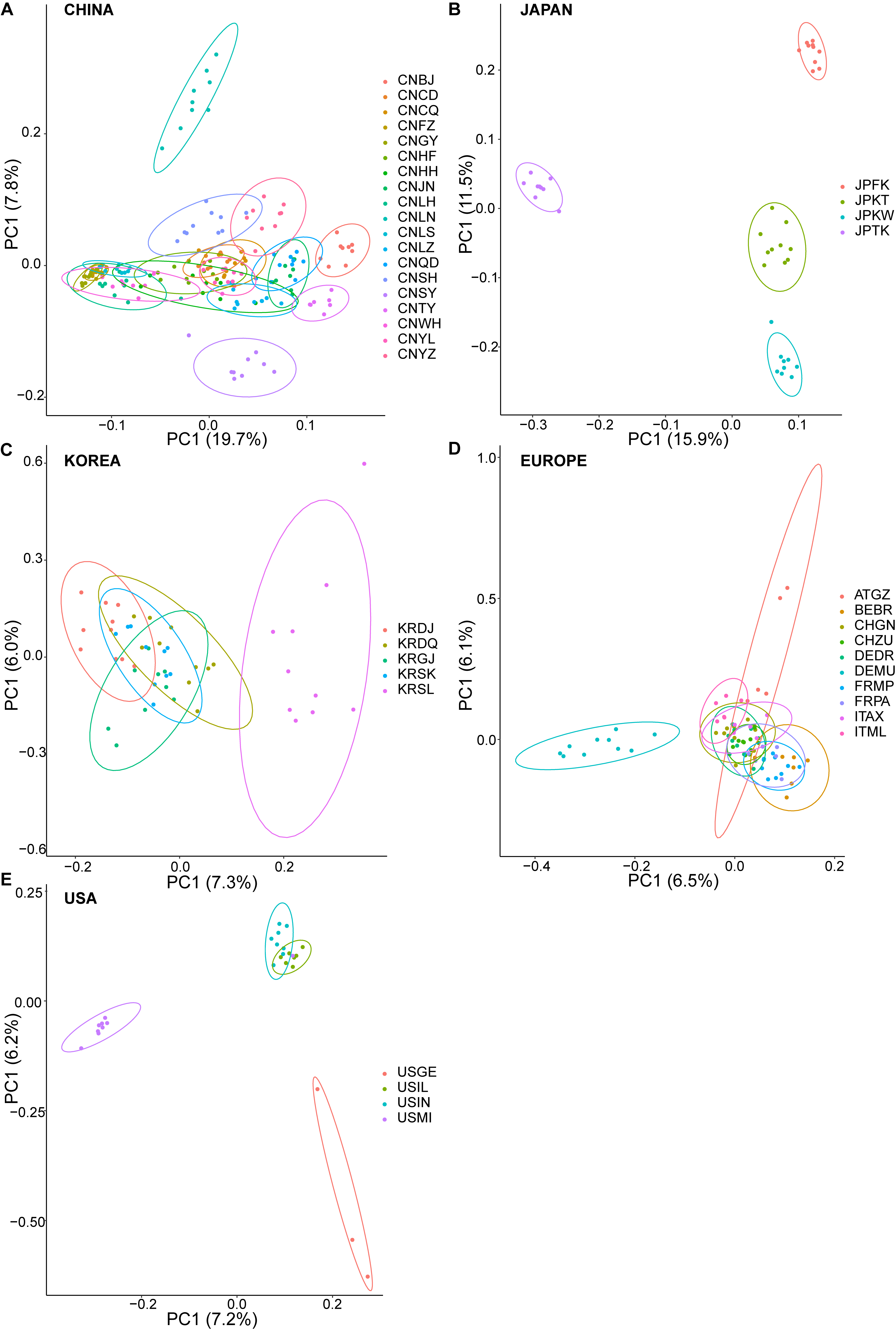

**Figure S7 Principal component analysis (PCA) plots based on genomic single nucleotide polymorphisms (SNPs) of samples from five different regions** All panels display PC1 plotted against PC2.

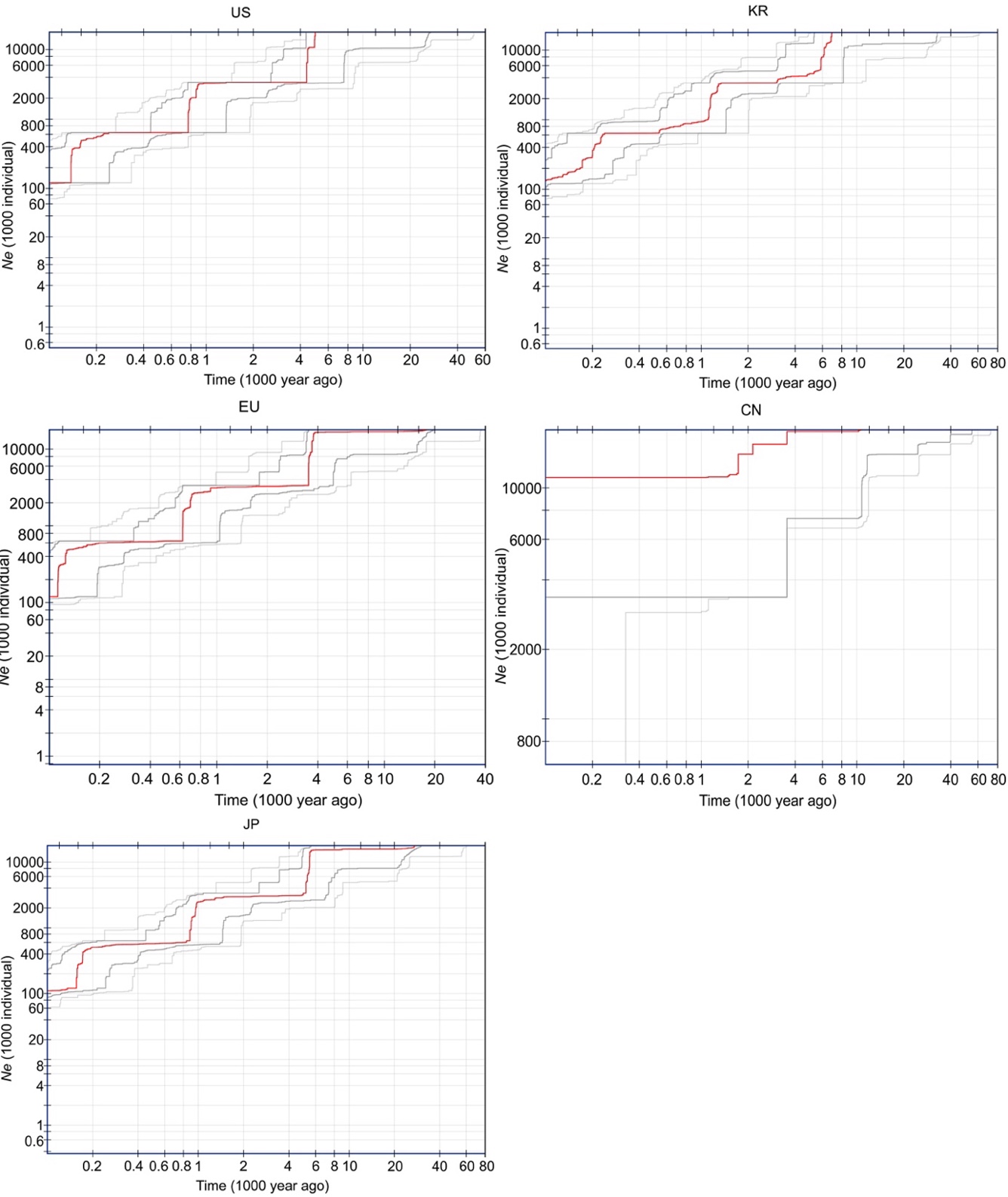

**Figure S8. Stairway plots depicting demographic history of populations from different regions.** Median estimates from 200 bootstraps are shown in red lines, while dark gray lines represent the 75% confidence interval and light gray lines represent the 95% confidence interval of the inference. Abbreviations: CN for China, EU for Europe, JP for Japan, KR for South Korea, and US for the United States.

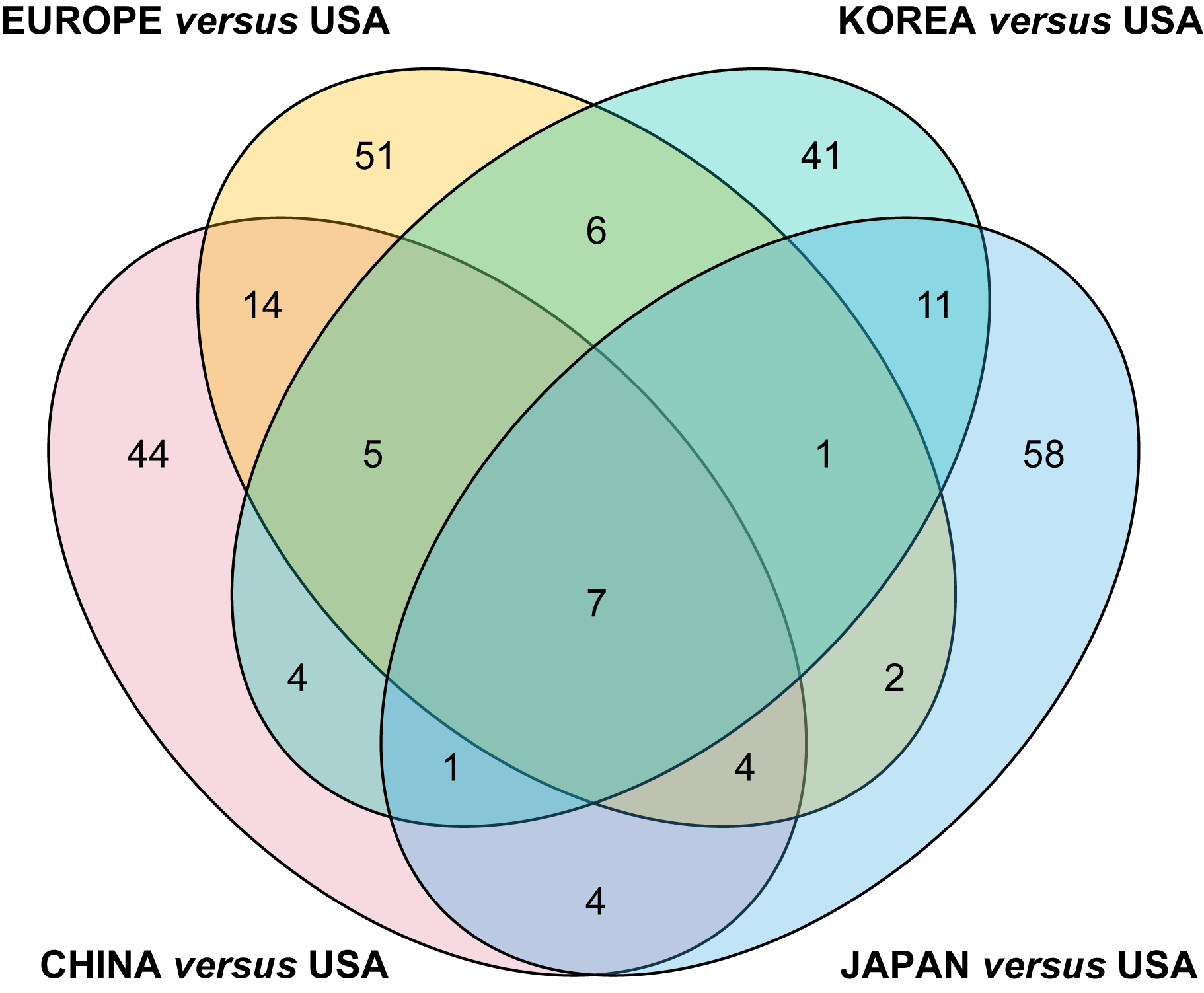

**Figure S9. Venn diagram illustrating overlapping genes identified in four pairs of invasive and native populations, each comprising 20 samples.** Please refer to Table S18 for the detailed list of samples included in this analysis.

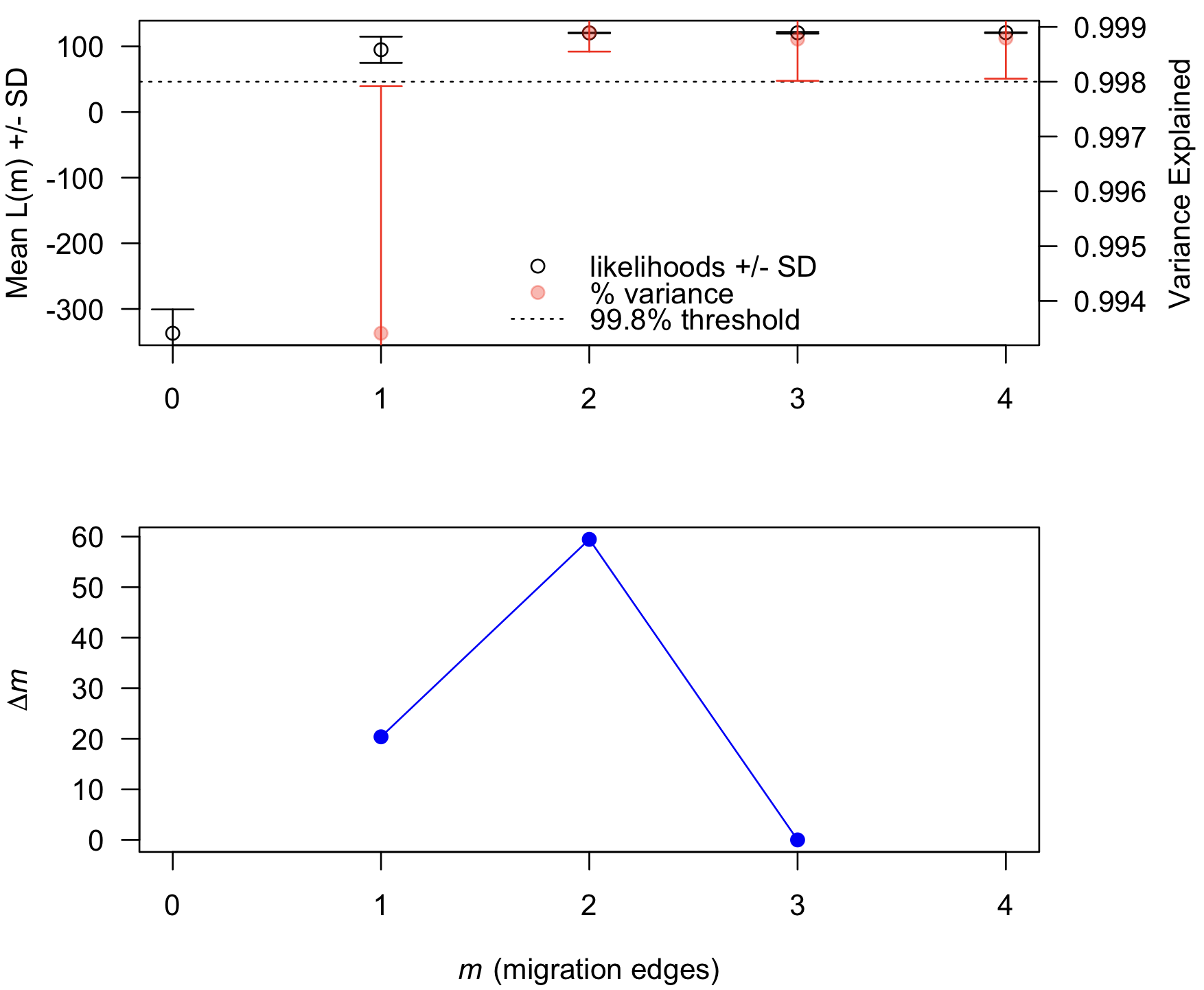
 **Figure S10. Testing of migration edges in the TreeMix analysis.** Two migrations (m = 2) are supported, as they account for 99.8% of the variances and yield the highest *Δm* value.
